# Structural Spine Plasticity in Olfaction: Memory and Forgetting, Enhanced vs. Reduced Discriminability after Learning

**DOI:** 10.1101/2020.12.04.411629

**Authors:** John Hongyu Meng, Hermann Riecke

## Abstract

How animals learn to discriminate between different sensory stimuli is an intriguing question. An important, common step towards discrimination is the enhancement of differences between the representations of relevant stimuli. This can be part of the learning process. In rodents, the olfactory bulb, which is known to contribute to this pattern separation, exhibits extensive structural synaptic plasticity even in adult animals: reciprocal connections between excitatory mitral cells and inhibitory granule cells are persistently formed and eliminated, correlated with mitral cell and granule cell activity. Here we present a Hebbian-type model for this plasticity. It captures the experimental observation that the same learning protocol that enhanced the discriminability of similar stimuli actually reduced that of dissimilar stimuli. The model predicts that the learned bulbar network structure is remembered across training with additional stimuli, unless the new stimuli interfere with the representations of previously learned ones.

## 1 Introduction

Structural reorganization of neuronal connectivity is thought to contribute to the mechanisms supporting learning and memory. The rates at which spines, which constitute the postsynaptic component of excitatory synapses, are formed and removed varies across brain areas. In most cortical areas, the rates for the formation and removal of spines are quite low in adult animals, and the overwhelming majority of spines (> 90%) are stable over many months, providing long-term memory for learned tasks [1–3].

In other areas, the spine dynamics are more pronounced. In the motor cortex, for instance, synaptic connections can exhibit an increased turnover rate *in vivo* within hours of training with a novel motor skill [4], and the stabilization of specific newly formed spines correlates with memory function [4, 5]. These newly formed spines are causally involved in learning the task, since shrinking them compromises the animal’s performance, while shrinking other spines in the same area does not [6].

The olfactory bulb, which is the first brain region to receive odor information, exhibits extensive structural plasticity. It involves the adult neurogenesis of granule cells (GCs), which are inhibitory interneurons, as well as the reciprocal synapses between GCs and mitral/tufted cells (MCs), which are the bulb’s principal cells. Compared to cortical areas, the dynamics of the spines of these synapses are substantially more pronounced; even over just a two-day interval, about 20% of the spines are removed and a comparable number of new spines are formed [7]. The olfactory system is therefore an excellent model to study structural plasticity.

Previous experiments have shown that on a time scale of days odor exposure enhances the stability of the spines in areas of the bulb that are activated by the odors [8]. Very recently, a more direct correlation has been demonstrated between the stability of individual spines and the activity of the GC that they are activating [9]. On shorter timescales (10 minutes), filopodia, which can be precursors of spines, appear and disappear [10]. Their dynamics have been found to depend on N-methyl-D-aspartate (NMDA) receptor activation on the GC dendrites [10], suggesting that the dynamics of the filopodia depend on the GC activity. Filopodia are also formed on spine heads. Their lifetime and orientation have been found to indicate the subsequent amplitude and direction of spine displacements, respectively, with the latter being modified by MC stimulation [11]. These combined contributions of GC- and MC-activity to the dynamics of filopodia suggest that the structural spine plasticity in the olfactory bulb may have a Hebbian character.

Since the structural plasticity in the olfactory bulb is an experience-dependent process, we wondered whether it contributes to the learning. Indeed, the perceptual learning of spontaneous odor discrimination involves the participation of adult-born GCs [12]. Modeling of adult neurogenesis suggests that this is due to a reduction in the similarity of the bulbar representations of those odors [13, 14]. Such a pattern separation was indeed observed as a result of active learning [15–17], where it also involved adult-born GCs [17], and after passive odor exposure [16]. Based on these observations, it is very surprising that odor discrimination training can also *increase* the similarities of the representations of those very training odors, i.e. training can reduce the discriminability of the training odors. In [16] this was observed when the training odors were very different from each other. For very similar odors, the same training protocol led to the expected decrease in similarity. This opposite behavior for dissimilar and similar odors can be thought of as balancing efficiency and robustness of odor perception in the learning process [16]. When training with very similar odors, it was found that some MCs become very sensitive to the slight differences between the odors through differential inhibition and differential disinhibition. However, the mechanisms behind these rich dynamics are still unclear.

In addition to the structural plasticity comprised of adult neurogenesis and rewiring of spines, synaptic-weight plasticity may also contribute to learning in the bulb [18, 19]. However, the nature of this plasticity is still poorly understood. Most of the common experimental protocols that trigger a consistent change in the synaptic weight in other brain regions lead to potentiation in some of the bulbar synapses and depression in others [19]. We therefore limit ourselves here to the structural plasticity. Motivated by the experimentally observed correlation between spine stability and GC activity [9], we focus on the spine dynamics and omit for now the impact of adult neurogenesis.

Here we show how a Hebbian-type model of structural plasticity can provide insight into the varied experimentally observed changes in MC responses during active and passive learning [16]. Our model addresses the dilemma between plasticity, which is required for learning, and stability, which is required for memory retention [20]. It retains the memory of a previous olfactory task when learning a new task as long as the bulbar representations of the odors defining the two tasks are sufficiently different to not interfere with each other. The model incorporates competition between different synapses on the same GC, which can be implemented by a shared limited resource.

## 2 Results

### 2.1 Formulation of the model

To investigate ramifications of the plasticity of the reciprocal synapses between MCs and GCs we focus on those two cell types and do not include the processing in the glomerular layer. Instead we consider the glomerular activation patterns as inputs to the MCs (Figure 1A). The MCs and GCs are described by firing-rate models governed by

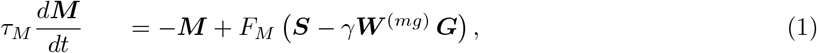

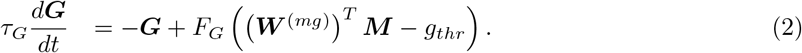

**Figure 1:**
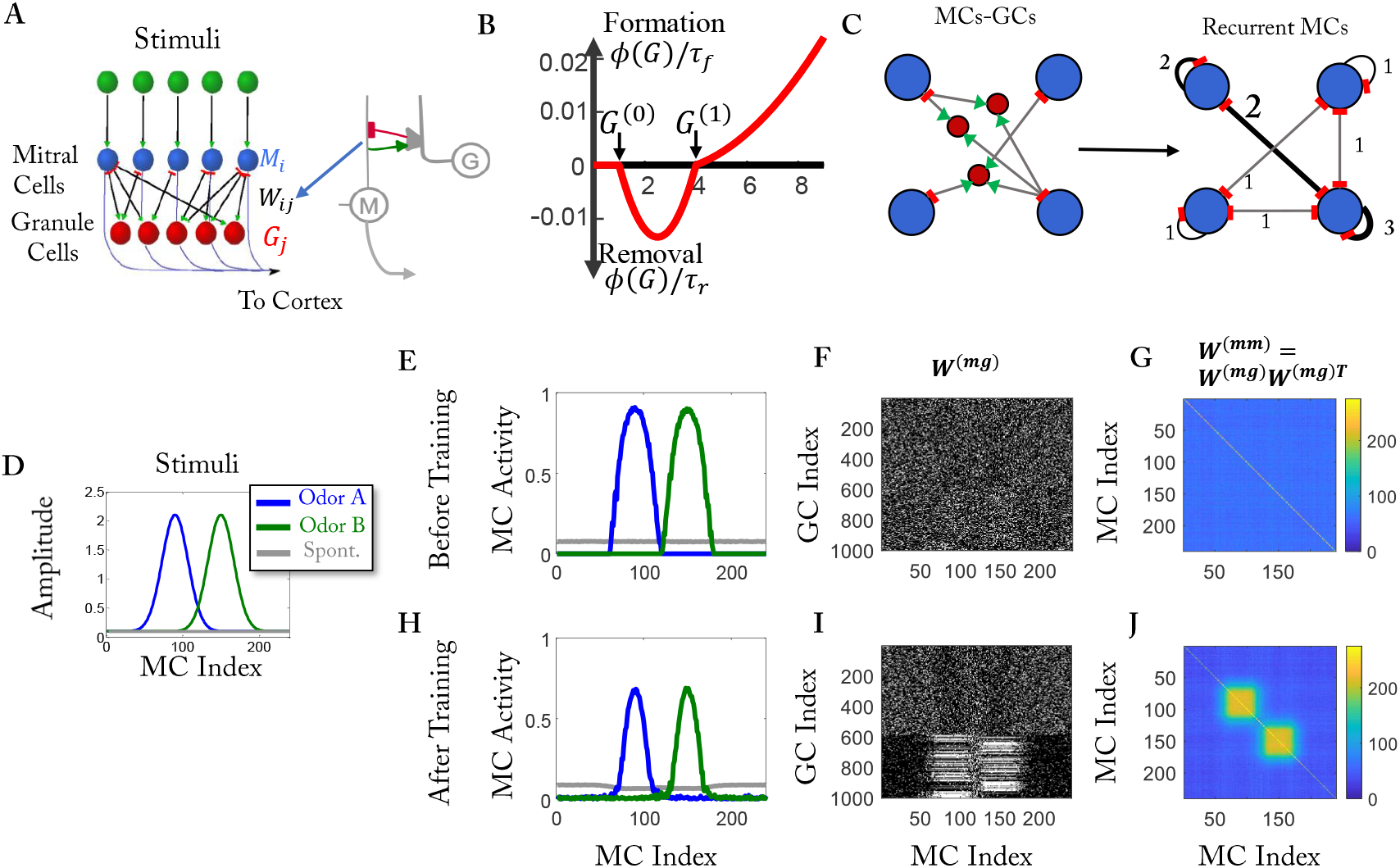
Computational model. (A) Sketch of the network. The synapses between excitatory MCs and inhibitory GCs are reciprocal. (B) Spine formation is controlled by *Mϕ*(*G*). Plotted is *ϕ/τ*_*f*_ for *ϕ* > 0 and *ϕ/τ*_*r*_ for *ϕ* < 0 (cf. Eq. 3) (C) Effective disynaptic recurrent inhibition of MCs. The numbers indicate the strength of the effective inhibition. (D) Simplified training stimuli. (E, H) MC activity before and after training, respectively. (F, I) Connectivity between MCs and GCs before and after training, respectively. Each white dot represents a connection between an MC and a GC. (G, J) Effective recurrent connectivity. The color represents the number of GCs that mediate the mutual inhibition of MCs (cf. C).

Here ***M*** and ***G*** are vectors of size *N*_*MC*_ and *N*_*GC*_, respectively, representing the firing rates of *N*_*MC*_ MCs and *N*_*GC*_ GCs. They are related to the neurons’ inputs via the activation functions *F*_*M*_ and *F*_*G*_, which are sigmoidal and piecewise linear, respectively (See Methods). The MCs receive sensory inputs ***S*** from the glomerular layer, and inhibitory input from GCs, with *γ* denoting the inhibitory strength. The GCs receive excitation from MCs, with *g*_*thr*_ > 0 setting a non-zero firing threshold for the GCs. Reflecting the reciprocal nature of the MC-GC synapses the connectivity matrix ***W*** ^(*gm*)^ is given by the transpose of ***W*** ^(*mg*)^.

Given the somewhat unclear nature of the synaptic-weight plasticity of the MC-GC synapses [18,19], all connections are assumed to have the same weight and ***W*** ^(*mg*)^ is taken to be either 1 or 0. Since the synapses are located on the secondary dendrites of the MCs, which reach across large portions of the olfactory bulb [21], we allow synapses to be formed between all MCs and all GCs without any spatial limitations. Thus, the ordering of the MCs and the GCs is arbitrary.

We focus here on the structural plasticity of the synapses. Our modeling of that plasticity is qualitatively motivated by a number of experimental studies. It has been shown that the overall stability of spines is enhanced if animals experience an enriched odor environment [8]. More precisely, across multiple days, spines in bulbar areas that were activated by the enrichment odors were stabilized, while those in other areas were not. Very recently, a positive correlation between the stability of a spine and the GC-activity of the dendrite on which it is located has been observed [9]. Since filopodia often are precursors of spines, we also glean information from experiments addressing their dynamics. The frequency of formation and removal of filopodia has been found to increase with NMDA-receptor activation [10] and to decrease with Mg^2+^-concentration. This suggests that the formation rate of spines increases with the activity of the GC-dendrite on which they are forming. The dynamics of filopodia that are located on spine heads are often indicative of subsequent dynamics of that spine. More specifically, the amplitude and direction of the movement of the spine head has been found to be correlated with the lifetime and orientation of such a filopodium, respectively [11]. Moreover, the dynamics of the filopodia have been found to be triggered by presynaptic glutamate release, suggesting a role of MC-activity in the spine dynamics [11]. Biophysical details about the mechanisms relating the spine dynamics with the temporal evolution of the neuronal activity during odor exposure are not well known yet.

Based on these observations, we assume the rate of formation and removal of a synapse connecting MC *i* and GC *j* to depend on the activities of those neurons. As a proxy for those activities we take the steady-state solution of Eqs.(1,2) for a given stimulus **S**. Inspired by the BCM model for synaptic weight plasticity [22], we express the formation and removal rates in terms of a single rate function,

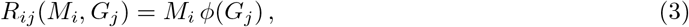

where *M*_*i*_ and *G*_*j*_ are the respective steady-state activities. The activation function *ϕ*(*G*_*j*_),

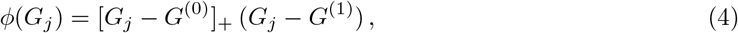

changes sign at a threshold *G*^(1)^ (Figure 1B), which controls whether a new synapse is formed or an existing synapse is removed (Figure 1B), giving the structural plasticity a bidirectional dependence on the GC activity. In addition, there is a second threshold, *G*^(0)^, below which the structural change is negligible. Specifically, in each trial of duration Δ*t*, which is one step in our simulation, a new synapse is formed with probability 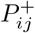,

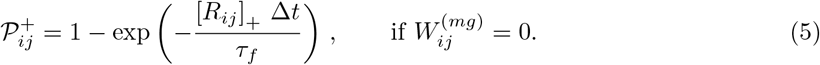

Conversely, an existing synapse is removed with probability 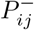

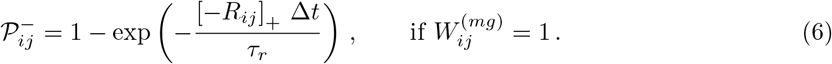

Here, *τ*_*f*_ and *τ*_*r*_ are the formation and removal timescale, respectively.

Since the change in the total number of synapses appears to be limited in experiments [23, 24], we hypothesize that there exists a homeostatic mechanism that keeps the total number of synapses on a given GC within a limited range. For most of our results, we employ a top-*k* competition mechanism, in which only the *k* strongest synapses survive. We then show that qualitatively the same results can be obtained with a more biophysical mechanism that is based on a limited resource (see Methods). Both mechanisms operate on a relatively fast time scale, as is generally needed for the stability of networks with Hebbian plasticity [25].

Since we are particularly interested in the impact of the plasticity on the ability of the network to discriminate between stimuli, we train it by exposing it to a random sequence of two stimuli A and B as sensory inputs ***S***. To illustrate the mechanism, we use simplified stimuli (Figure 1D), while naturalistic stimuli are used to compare with experimental data (Figure 2B).

**Figure 2:**
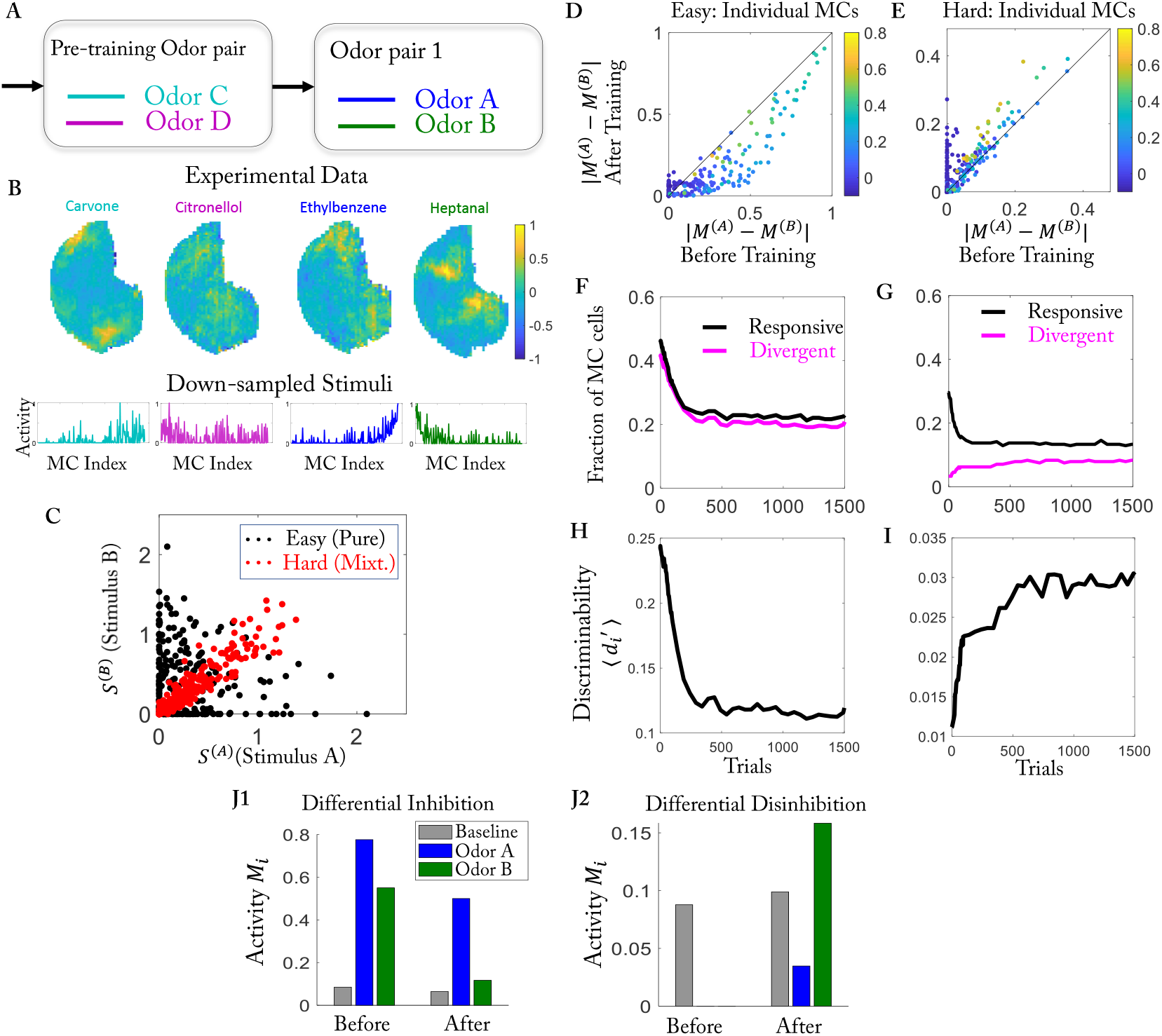
Easy and hard discrimination task using naturalistic stimuli. (A) Training protocol. (B) (Top) Glomerular activation patterns (cf. [27]). (Bottom) Activation patterns down-sampled to 240 points, serving as stimuli **S**^(*i*)^. MCs are sorted based on the difference in activation by the stimuli of odor pair 1. (C) Activity of each MC for the two stimuli used in the easy (black) and hard (red) task. (D) Difference in response for each MC before and after training for the easy task. Color represents the mean response ***M*** ^(1)^ + ***M*** ^(2)^ 2***M*** ^(*air*)^ */*2 before the training. (E) As (D), but for the hard task. (F, G) Temporal evolution of the number of responsive MCs and divergent MCs during the training in the easy task and hard task, respectively (cf. [16]). (H,I) Temporal evolution of 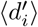 for the easy task and the hard task, respectively. (J) Training for the hard task enhances differences in responses of the MCs to the two odors. (J1) This MC changes from being excited by both odors to being excited by only one of them. (J2) This MC changes from being inhibited by both odors to being inhibited by only one of them. Parameters as in Table 1 except for *γ* = 1.7e−4.

Summarizing the algorithm, in each step, the steady-state firing rates ***M*** and ***G*** in response to a randomly chosen odor ***S*** from the training set are computed. Based on the resulting formation/removal rates *R*_*ij*_ the connectivity matrix ***W*** ^(*mg*)^ is updated in two steps: first, implementing homeostasis by top-*k* competition, on each GC only the *k* synapses with largest *R*_*ij*_ are retained. Second, in the learning step the connectivity is updated based on the Hebbian learning rules (5, 6). The homeostatic step always precedes the learning step to avoid potential artificial oscillations. We simulate the model long enough for the connectivity to reach a statistically steady state as measured by the discriminability of the training odors (see Methods for details). As initial condition, we take a connectivity in which each GC is randomly connected to *N*_*conn*_ MCs. Unless specified otherwise, the model parameters are as listed in Table 1. The steady states are obtained by setting *τ*_*G*_ = 0, solving Eq.(1) using ODE45 in Matlab, with ***G*** solved from Eq.(2).

**Table 1:** Parameters used in the simulations of the model (unless stated otherwise).

|  |  |  |  |
| --- | --- | --- | --- |
| $N_{MC}$ | 240 | $N_{GC}$ | 1000 |
| $N_{conn}$ | 60 | k | 66 |
| $\tau_M$ | 1 | $\tau_G$ | 0 |
| $\Delta t$ | 1 | | |
| $g_{thr}$ | 4.4 | $\gamma$ | 5e-4 |
| $G^{(0)}$ | 1 | $G^{(1)}$ | 4 |
| $1/\tau_f$ | 6e-4 | $1/\tau_r$ | 6e-3 |

### 2.2 Training the model with a pair of simplified stimuli

To illustrate the network evolution resulting from this plasticity rule, we use a pair of simple model stimuli A and B that excite partially overlapping sets of MCs (Figure 1D). In the absence of any odors (‘air’) the input is taken to be non-zero and homogeneous, reflecting the spontaneous activity of the MCs or of the neurons driving their input. Initially, the MC-GC connectivity is random (Figure 1F), resulting in the MC activities shown in Figure 1E. After training, many GCs are preferentially connected with MCs that are activated by the stimuli (Figure 1I). Due to the reciprocal nature of these synapses, this induces enhanced mutual disynaptic inhibition between co-activated MCs as seen in the effective connectivity matrix *W* ^(*mm*)^ = *W* ^(*mg*)^*W* ^(*mg*)*T*^ (Figure 1G, J). It gives the number of GCs that mediate the inhibition between the respective MCs and reflects the stimulus patterns used in the training.

The learned network structure reflects the effectively Hebbian character of the structural plasticity rule (3) through which connections are established between cells that fire together. If a GC responds to a stimulus by integrating excitatory inputs from activated MCs, other MCs that are activated but not yet connected to this GC have a large probability of forming a connection with that GC. As a result, if two MCs respond to the same stimulus, the number of GCs they both connect to increases if the model is trained by that stimulus. This is reflected by the two blocks along the diagonal line of the effective connectivity matrix (Figure 1J). Since the synapses are reciprocal, MCs that drive more GCs also receive more inhibition. As a result, the training preferentially reduces the activity of initially highly activated MCs (Figure 1E, H), consistent with observations [12, 26].

The overall behavior of the model is robust with respect to changes in its parameters (Supplementary Materials).

### 2.3 Training enhances or reduces discriminability depending on stimulus similarity

We used our model to investigate the intriguing result of [16] who found that the learning process enhanced the discriminability of stimuli *only* if they were very similar, i.e., for hard discrimination tasks. If the stimuli were dissimilar, i.e., if the task was easy, the learning actually reduced their discriminability. We used the same training protocol as in the experiment and use naturalistic stimuli as inputs, adapted from glomerular activation data of the Leon lab [27] (Figure 2B (top)) and down-sampled to reduce the computational effort (Figure 2B (bottom)). These stimuli were used for the easy discrimination task. As in [16], we used mixtures of the same stimuli for the hard discrimination task (60% : 40% *vs.* 40% : 60%, see Methods). The two odors of the hard task drove each MC to a very similar degree (Figure 2C).

Our model successfully reproduced several experimental observations in [16]. First, the difference in the response of the MCs to the two odors in the odor pair *decreased* for the majority of the MCs after training in the easy task, while it *increased* in the hard task (Figure 2D, E). Here the response was defined as the difference between odor-evoked activity and air-evoked activity. Especially in the hard task, a fraction of cells showed no significant response difference before training, but an essential response difference after training (Figure 2E, dots along the y-axis). As in [16], we quantified these changes classifying MCs into responsive cells, i.e., cells that showed a significant response to at least one of the two odors, and divergent cells, for which the two odors evoked significantly different responses (see Methods). In the easy task, the number of divergent cells decreased along with the number of responsive cells (Figure 2F). In contrast, in the hard task, the number of divergent cells increased by a small amount, although the number of responsive cells still decreased (Figure 2G). The results were not sensitive to the threshold *θ* that classified the cells (Figure S12). Second, we used 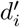 (see Methods) of each divergent neuron to measure the dissimilarity of the response patterns. With training, the mean 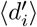 went down in the easy task (Figure 2H), indicating reduced discriminability, while it went up in the hard task (Figure 2I). Third, as observed in [16], in the hard task, some of the MCs became divergent by differential inhibition or by differential disinhibition. Thus, some MCs were excited by both odors before training, but only by one of them after training (Figure 2J1). Conversely, some other MCs were inhibited by both odors before training, but only by one of them after training (Figure 2J2). Figure S1 gives an overview of all MCs that showed such differential behavior.

### 2.4 Network connectivity underlying the reduction/enhancement of discriminability

To better understand the mechanism underlying the change in discriminability, we trained the network using pairs of simplified stimuli (Figure 3C). As in the case of realistic stimuli, the stimuli for the hard task were mixtures of two easily discriminated stimuli. Again, we first exposed the network to a set of pre-training stimuli that set up an initial connectivity that was independent of the task stimuli (Figure 3A). It affected the initial response of the network to the training stimuli (Figure 3E, I). Thus, at the beginning of the training, the MCs for which the difference in the input from the two training stimuli was maximal were not necessarily the MCs that exhibited the maximal difference in their response to those training stimuli (gray and black arrows, Figure 3H, L).

**Figure 3:**
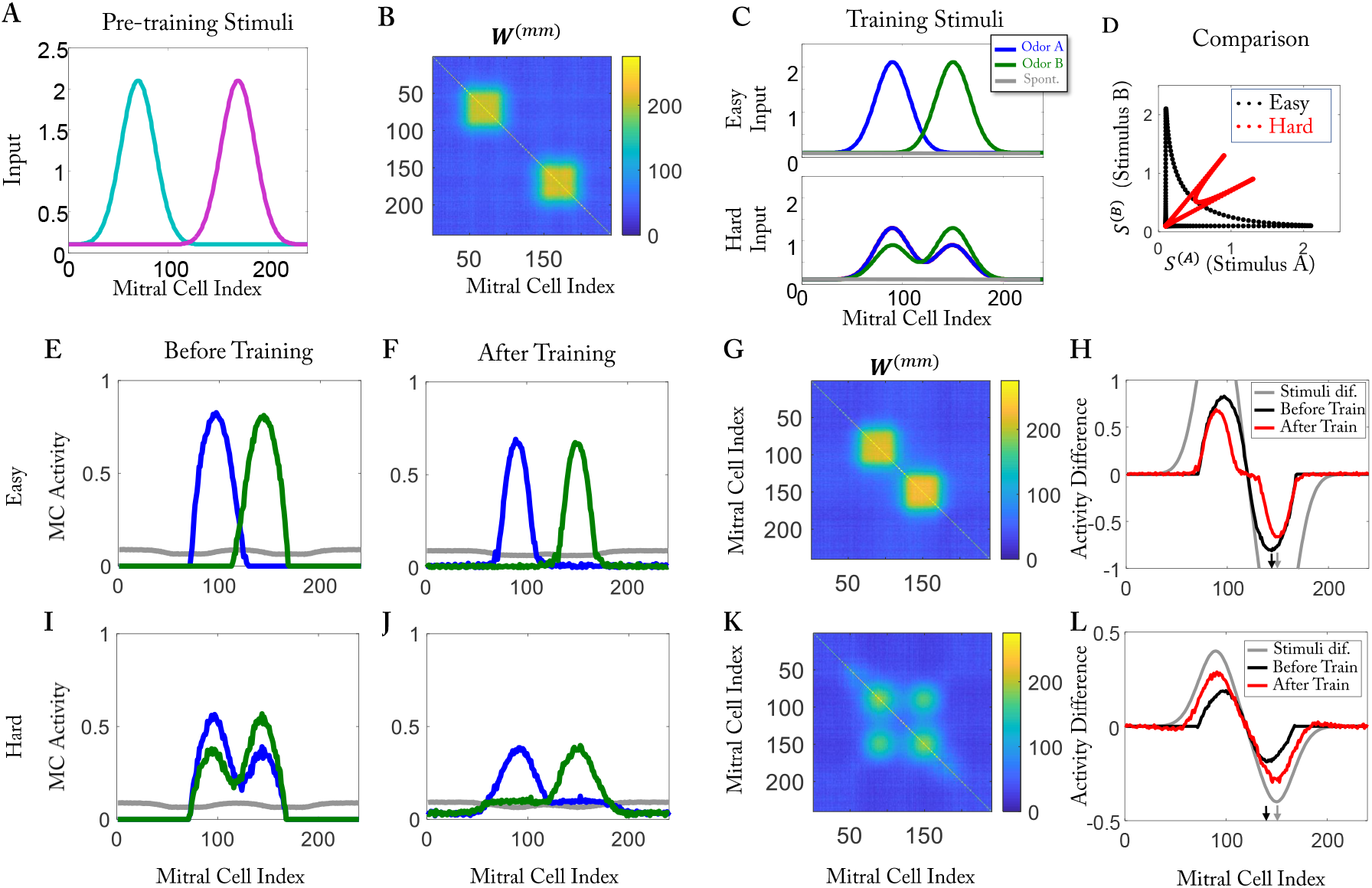
Training with simplified stimuli. (A) Stimuli for pre-training. (B) Effective connective matrix after pre-training. (C) Training stimuli for the easy task (top) and the hard task (bottom). (D) Activity of each MC for the two stimuli used in the easy (black) and hard (red) task. (E-H) Easy task: (E) MC activity before training (after pre-training). (F) MC activity after training. (G) Effective connectivity matrix after training. (H) Activity difference before (black) and after training (red). The grey line gives the difference in the inputs (cf. (C)). (I to L) As (E-F) but for hard task: in (H, L) the arrows indicate the maximum of the difference in the stimuli (grey) and in the MC activity (black). In (H), the MC for which the difference in the training stimuli is maximal is the same as in (L).

The repeated exposure of the training stimuli modified the connectivity and the MC responses. In the easy task, the effective connectivity had only two blocks after training, which were along the diagonal and corresponded to odors A and B, respectively (Figure 3G). This inhibition reduced the MC activity and with it the number of responsive cells. Since most MCs responded essentially to only one of the two stimuli, the response difference decreased along with the overall response (Figure 3H, red line vs. black line). In the hard task, both stimuli activated the same set of MCs, just to a different degree, reflecting the difference in the concentrations of the two components. The discriminability of the stimuli was therefore not compromised if the MC activities were reduced, as long as the difference in the representations of the two stimuli were sufficiently maintained. The effective connectivity resulting from the training achieved this. It had two additional, off-diagonal blocks (Figure 3K). While in the easy task each activated MC only received disynaptic inhibition from MCs that are activated by the same odorant, in the hard task each activated MC received disynaptic inhibition from MCs that were activated by either of the two components of the mixture. Even though the effective inhibition from the MCs activated by the same mixture component increased with the concentration of that component, which reduced the response difference, the effective inhibition from the MCs activated by the other component decreased with the concentration of that first component. This compensation acted to preserve the response difference. In fact, for a large number of MCs the difference in the response even increased with training (Figure 3J, L). Assuming that for a cell to be classified as divergent, this difference had to be above some threshold, the number of divergent cells decreased in the easy task, while it increased in the hard task (Figure 3H, L, cf. Figure S12), as we observed for the naturalistic stimuli (Figure 2F, G). Qualitatively similar results were obtained without a pre-training phase (Figure S9).

Note that d-prime depends on the difference in the response relative to the response variability. Since we assume that the firing rates of our model correspond to spike trains with a Poisson-like variability that increases with its mean, a reduction in the mean firing rates implies a reduction of the variability. As a consequence, a reduction of the mean that essentially preserves the difference as seen in Figure 3J, L enhances d-prime (Fig.2I).

### 2.5 Forgetting of connectivity induced by training with interfering stimuli

The qualitative agreement of the model with the experimental findings suggests that the olfactory bulb stores the memory of previous tasks in its connectivity. We therefore asked to what extent this memory is affected by subsequent exposures to new stimuli. Could they compromise previous memories and diminish the discriminability of previously learned stimuli? Would the system forget learned tasks? To address this, we trained our network sequentially in three phases (Figure 4A). In the first and second phases, we used two different, hard tasks. In the third phase, the network was again trained on the same task as in the first phase.

**Figure 4:**
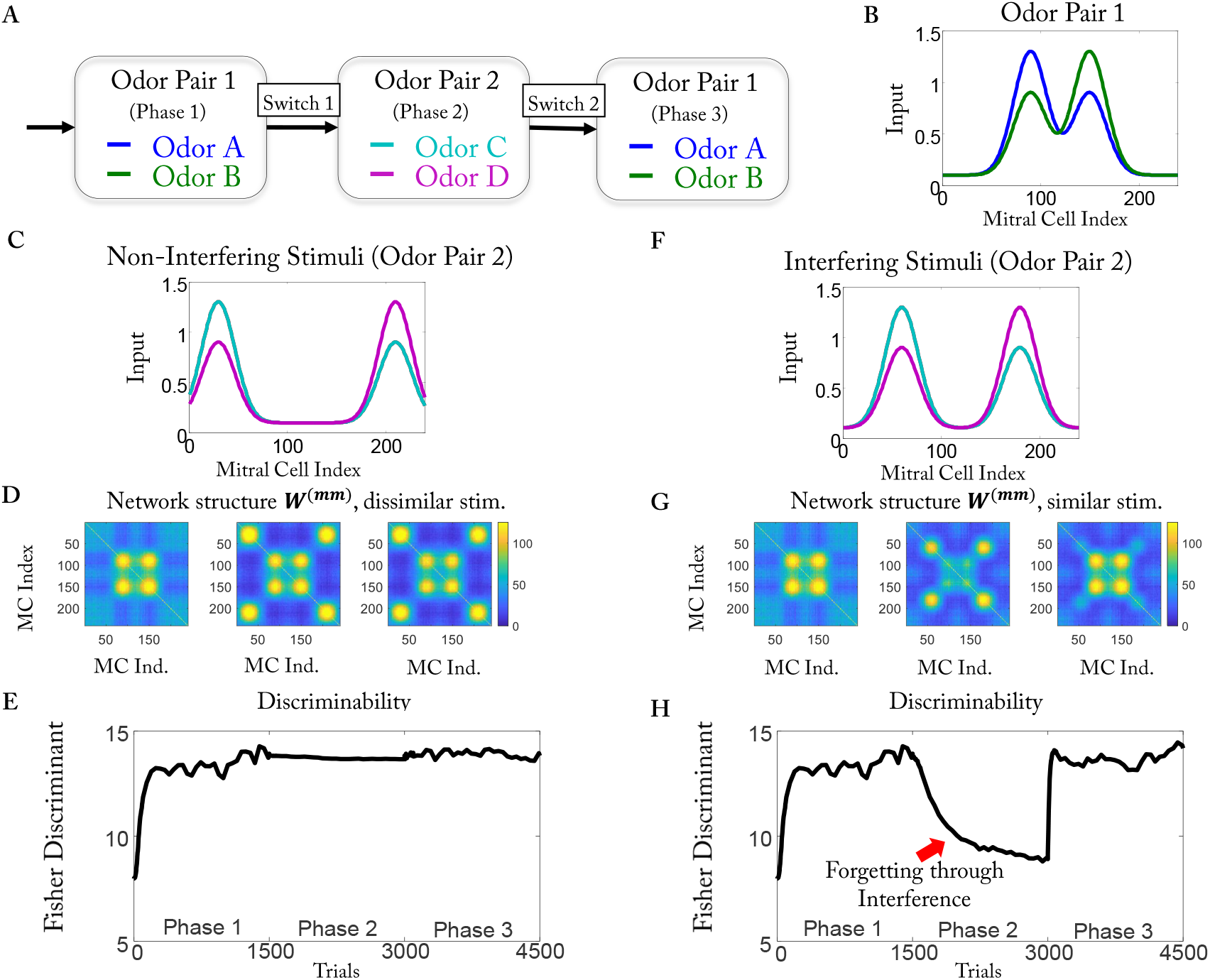
Interference leads to forgetting. (A) Training protocol. (B) Stimuli of odor pair 1. (C, D, E) Non-interfering stimuli. (C) Stimuli of odor pair 2. (D) Effective network structure *W* ^(*mm*)^ at the end of the training phases. The network remembers the previously trained structure. (E) Fisher discriminant of odor pair 1 as a function of training time. Performance is not impacted by training with pair 2. (F, G, H) As (C, D, E) but for interfering stimuli. (G) The network forgets most of the previously learned structure. (H) The Fisher discriminant of pair 1 decreases during phase 2: interference introduces forgetting. After re-training with odor pair 1 in phase 3, the Fisher discriminant returns to the original value: learning ability is intact.

The behavior of the model depended strongly on the similarity of the two odor pairs (Figure 4B, C, F). When the MCs that were activated by the stimuli of odor pair 2 did not overlap with those of odor pair 1, the network preserved the previously learned structure (Figure 4D). However, if there was significant overlap, the previously learned structure was forgotten during phase 2 (Figure 4G, middle, panel). In the following discussion, we will refer to the stimuli in Figure 4C, F as non-interfering and interfering stimuli, respectively. The previously learned structure was re-learned by re-training in phase 3 (Figure 4G, right panel). To quantify the evolution of the discriminability, we used the Fisher discriminant 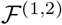, which is given by the sum over the squares of the 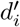 of the individual MCs in response to stimulus pair 1 and 2, respectively (see Methods). The Fisher discriminant 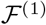 remained high during the training with odor pair 2 if that pair was non-interfering (Figure 4E), while it decreased if odor pair 2 was interfering (Figure 4H). We expect that animals will spontaneously discriminate the odors in a pair only if their bulbar representations are sufficiently different, i.e., if 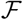 is sufficiently large [12]. Thus, the model predicts that at the end of phase 2 of a multi-phase perceptual learning task using consecutive, strongly interfering odor pairs, an animal will not spontaneously discriminate any more the odors it learned in phase 1, because the training in phase 2 reduces the difference in the bulbar odor representations of the odors of pair 1. However, for sufficiently different odor pairs, the spontaneous discrimination should remain intact. In addition, the model predicts that re-learning in phase 3 is faster than the initial learning in phase 1 and forgetting is slower than either of the learning processes. Note that repeatedly switching between odor pairs did not impair the learning ability (Figure S10).

### 2.6 Realization of competition through competition for a limited resource

In order to keep a Hebbian model stable, different compensatory processes can be imposed depending on the state of nearby synapses on the same dendrite [25]. In the model, we have discussed so far, we used top-*k* competition because of its simplicity, without identifying a specific, biologically feasible mechanism. In this section, we discuss one such mechanism.

The idea behind the competition is that within a single neuron, possibly within a single dendritic branch, resources used for the formation of synapses are limited. In hippocampal CA1 neurons, stimulating spines leads to the shrinking of neighboring unstimulated spines [28]. A plausible explanation of this phenomenon is that some resource is limited within the neuron. This idea has been implemented in a computational model for synaptic-weight plasticity [29]. Along similar lines, we assumed that the formation of each synapse depletes a resource pool by a fixed amount, which is returned to the pool if the synapse is removed at a later time. We took the activation function *ϕ*(*G*) in (3) to depend on the fill-level *P* of that resource pool (Figure 5A, red curve) and assumed that it is comprised of a formation term *ϕ*_*form*_ (Figure 5A, blue curve) and a removal term *ϕ*_*rem*_ (Figure 5A, green curve),

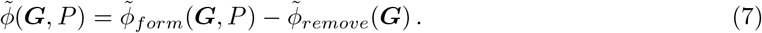

**Figure 5:**
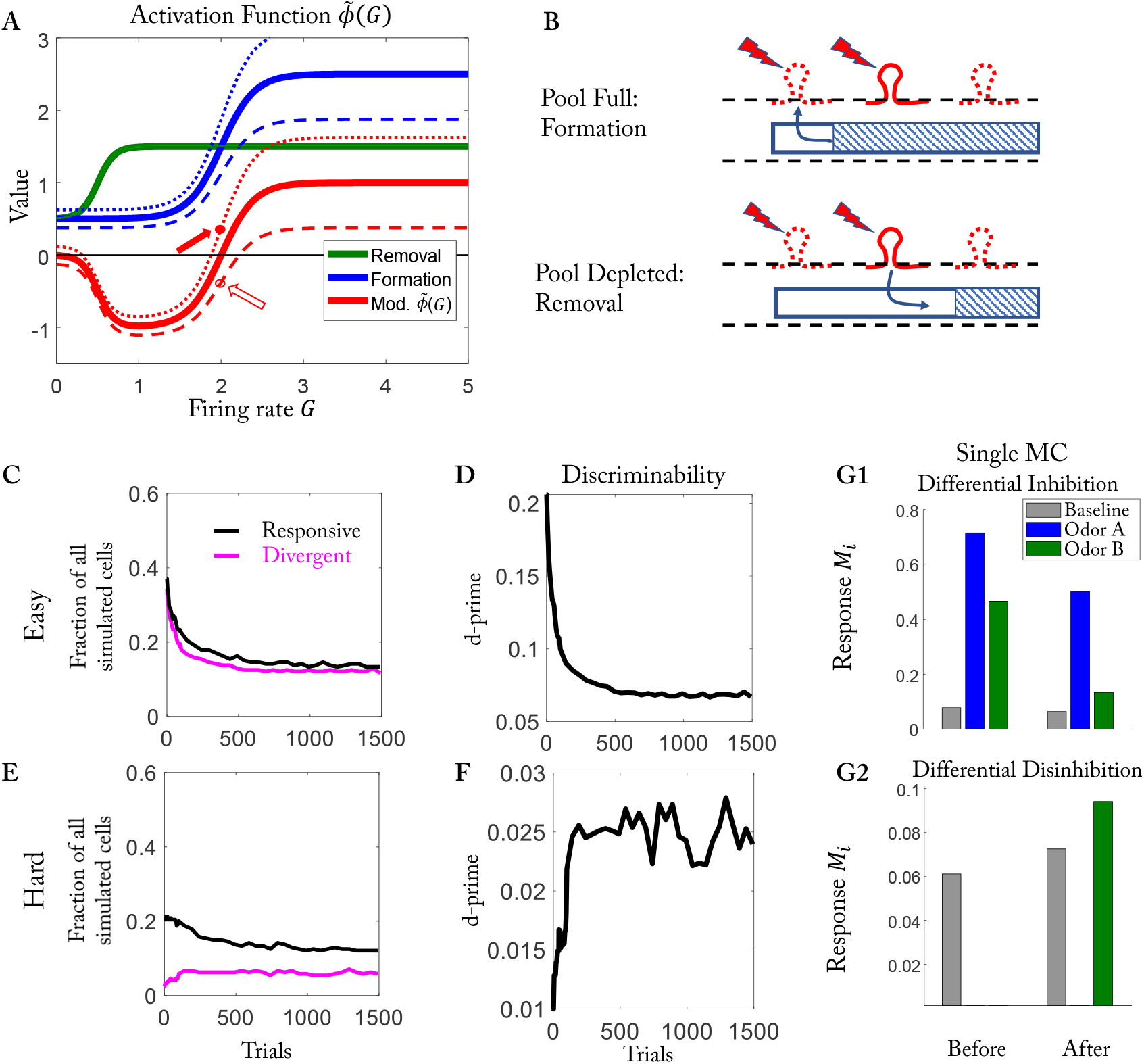
Realization of competition by a common resource pool. The training protocol is the same protocol as in Figure 2. (A) The modified activation function 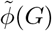 (red) is the difference between the formation function (blue) and the removal function (green). *P* = *P*_0_ (thick), *P* > *P*_0_ (dotted), *P* < *P*_0_ (dashed). (B) (Top) For *P* > *P*_0_ synapses tend to form (solid arrow in (A)). (Bottom) For *P* < *P*_0_ synapses tend to be removed even though the GC activity is the same (open arrow in (A)). (C to G) Results as in Figure 2F-J for the resource-pool model.

Both terms depended on the activity ***G*** and, in addition, *ϕ*_*form*_ depended on the current size of the resource pool,

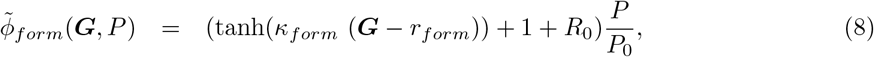

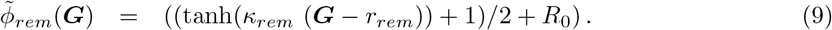

Here *P*_0_ is the equilibrium size of the resource pool for which no spines are formed or removed when the GC is not active. The fill-level *P* of the resource pool depended on the current number of synapses *n*, *P* = *P* ^(*all*)^ − *n*, where *P* ^(*all*)^ is the total amount of the resource in the cell, which was assumed to have the same constant value for all GCs. In analogy to the original model, we defined the two thresholds 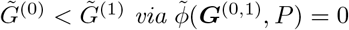.

It is easy to see how the limited resource pool provides a homeostatic mechanism. For very low GC activity and *P* < *P*_0_ one has 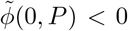 leading to the removal of synapses, which replenishes the pool until *P* reaches *P*_0_. Conversely, 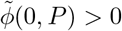 for *P* > *P*_0_, which leads to the formation of new synapses, decreasing *P* until it reaches *P*_0_. Thus, for low GC-activity *P* eventually goes to *P*_0_ and the number of the synapses on that GC goes to the initial value *n* = *N*_*conn*_ = *P* ^(*all*)^ − *P*_0_.

In [16], mice were exposed to the test odors only when they were on the training stage. When they were back in the cage, the training odors were absent, and the activities of GCs corresponding to those odors were likely to be low. In the original model, the activation function 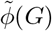 vanishes when *G* < *G*^(0)^, which means synapses are neither formed nor removed. Thus, the network is static. However, in the modified model, the activation function 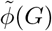 is in general non-zero when *G* < *G*^(0)^. As a result, periods of low input also lead to changes in the connectivity. Here we include such ‘air’ trials, during which only spontaneous activity is presented, after every four training trials. During air trials, the GC activity is very low and synapses are formed and removed based particularly on the size of the respective resource pool, pushing each pool towards *P*_0_. Given the uniform input to the MCs during air trials and the dependence of spine formation on MC activity (cf. (3)), the formation is biased towards MCs that receive less inhibition, i.e., that have fewer connections with GCs, while the removal is biased towards MCs that receive more inhibition. We find that inserting such air trials increases the dynamics of the synapses and speeds up the convergence of the network evolution.

The formation of synapses on significantly active GC is controlled by 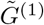. As more synapses are formed and the fill-level decreases, 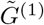 increases. This makes it more difficult for synapses to form on that GC, leading to a saturation of the number of its synapses. This dependence of the threshold 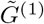 is reminiscent of the sliding threshold of the BCM model for synaptic-weight plasticity [22]. However, the sliding is caused here by the changes in the limited resource instead of a temporal average over the previous activities. The change in 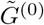 is opposite to the change in 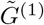 when the size of the resource pool changes.

It is worth mentioning that in this resource-based model, an existing synapse is removed only if the activation function is negative, 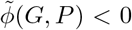 < 0, while in the top-*k* competition model, synapses can be removed not only when *ϕ*(*G*) < 0, but also directly by that competition.

With this biophysically plausible stabilization of the structural plasticity, our model yielded results that were qualitatively similar to those obtained with the simpler, original model (compare Figure 5C-G and Figure S11). The additional parameters we used in this section are listed in Table 2.

**Table 2:**
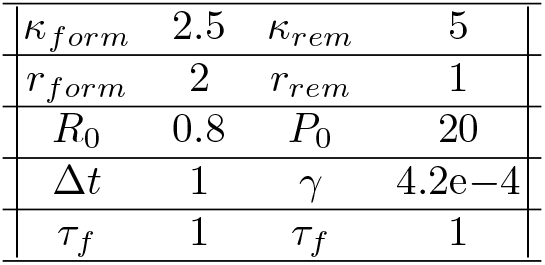
Table of parameters for the resource-pool model (unless stated otherwise).

## 3 Discussion

In the olfactory bulb, structural plasticity is operating on a timescale that is comparable to some learning processes. This raises the question of what role structural plasticity plays in these processes. In this study, we proposed a simple Hebbian-type model to address this question. Our model shows that a local unsupervised learning rule is sufficient to explain the intriguing observation that learning can have an opposite impact on the discriminability of odors for easy and for hard tasks [16]. While the discriminability of very similar odors is enhanced by the learning, it is reduced for dissimilar odors. In fact, for the dissimilar odors, the number of MCs that respond differently to the odors decreases with learning, deteriorating the performance. Moreover, the model captures the observation that through the learning, individual MCs can become very sensitive to small differences in the odor patterns *via* differential inhibition or differential disinhibition [16].

In contrast to a previous model for structural spine plasticity [7], this model describes a learning mechanism that allows the network to remember previously learned tasks in its connectivity. These memories are robust against subsequent learning of dissimilar tasks. The learned connectivity is only erased by exposure to interfering stimuli, which overlap significantly with previously learned ones. This prediction should be amenable to experimental tests using sequential perceptual learning [30] with two odor pairs, each comprised of two very similar odors. Repeated exposure with one of the two odors of the first pair will enhance the spontaneous discriminability of the odors in that pair. The subsequent repeated exposure with an odor from the second pair is expected not to affect the discriminability of the odors in the first pair if the odors in the second pair are dissimilar from the first pair. However, if there is a significant overlap between the pairs, the second training is expected to compromise the discriminability of the first pair. Note, however, that if the odors in the second pair are very similar to those in the first pair, this interference is not expected. The latter aspect would differentiate this forgetting from the widely observed retrieval-introduced forgetting [31].

We discussed two fast homeostatic mechanisms that stabilize the network activity despite the Hebbian-type learning rule: an abstract top-*k* competition and a biologically plausible competition of synapses for a limited resource. Changes in the availability of that resource lead to the sliding of a threshold for the formation of synapses, which is reminiscent of the BCM-model [22]. It is worth noting, however, that different brain areas may have different stabilization mechanisms, which limits the generality of that aspect of our model. Moreover, resource-dependent competition may be very local and may not lead to competition across different dendrites or the whole neuron. In addition, heterosynaptic plasticity may be signal dependent rather than resource-dependent [28]. For these reasons, we leave open the question of how to implement the stabilization mechanism.

A characteristic feature resulting from the structural plasticity implemented in our model is the emergence of subnetworks of excitatory and inhibitory cells that are specific to the learned stimuli. Due to the Hebbian nature of our model for spine formation, strongly connected MC-GC pairs share receptive fields, i.e., they exhibit selectivity for the same stimuli. Since the MC-GC synapses are reciprocal, GCs give strong inhibition to those MCs from which they receive many connections. This results in the mutual, disynaptic inhibition of MCs with similar receptive fields that underlies the enhancement of stimulus discrimination by learning.

Reciprocity at the level of individual synapses, as it is found in the olfactory bulb, is not common in the brain. Nevertheless, recent observations have revealed similar circuit structures in visual cortex V1 [32, 33] and in the posterior parietal cortex [34]. In V1, learning of a visual discrimination task enhanced the selectivity not only of pyramidal cells, but also of interneurons, with parvalbumin-positive interneurons (PV) becoming as selective as pyramidal cells [32]. Strikingly, this occurred although anatomically PV cells are connected with the majority of nearby pyramidal cells, which in mice have very different receptive fields. It was found, however, that the strengths of the connections between pyramidal cells and PV cells varied substantially between cell pairs and were co-tuned, such that PV cells give back strong inhibition to those pyramidal cells from which they receive strong excitation [33], yielding a structure not unlike that of the bulbar MC-GC network. The plasticity mechanism leading to this structure is not well understood yet; a modeling study has shown that a combination of excitatory and inhibitory plasticity can lead to this structure, if the two mechanisms synergistically complement each other [35]. Analogously, in the posterior parietal cortex, the selectivity of excitatory and inhibitory neurons has been found to be similar in terms of the animal’s choice during decision making [34].

While structural spine plasticity is a striking feature of the olfactory bulb, it is not limited to excitatory-inhibitory recurrent networks. The benefit of such plasticity has also been addressed for other network structures. Among feedforward networks, a two-layer network has been considered that is to infer its inputs from the activities of its output neurons [36]. There, Hebbian structural plasticity with a fixed magnitude of the connection weights was found to outperform that of Hebbian synaptic-weight plaxsticity for fixed connectivity structure, if the connectivity was sparse. The connectivity of the olfactory bulb is very sparse. Each of the about 5 million GCs connects with on the order of 50-100 of the 50,000 MCs [37]. Considering that MC dendrites can reach across large portions of the bulb [21,38], which reflects the lack of significant chemotopy and the high dimensionality of odor space, less than 1% of the possible connections are actually made. This makes structural plasticity extremely beneficial, since it provides a mechanism to focus the connections on the most relevant neurons.

Hopfield-like associative networks with synaptic-weight plasticity typically suffer catastrophic forgetting of memories when the number of learned memories exceeds the network capacity. Computational modeling has shown that structural plasticity can replace this by a graded deterioration of recently learned memories [39], without invoking the replay of previously learned memories or the explicit use of history-dependent terms (e.g. [40]. The analogous limitations of recurrent excitatory-inhibitory networks like that of the olfactory bulb with respect to the number of discriminable activity patterns and the type of degradation that occurs as the number of learned stimuli is increased has not been investigated yet in any detail.

In addition to spine plasticity, the olfactory bulb is characterized by substantial neurogenesis of GCs, even in adult animals. Functionally, the adult-born GCs (abGCs) contribute substantially to the animal’s ability to learn to discriminate very similar odors, both for perceptual learning [12,41] and for active learning [17]. Many experiments found that the survival of the abGCs depends on their activity [42, 43]. Modeling studies showed that such activity-dependent survival naturally leads to bulbar network structures that are quite similar to those resulting from structural spine plasticity discussed here and similarly enhance odor discrimination [13, 14]. Recent experiments have questioned, however, the previously reported extensive cell death of abGCs [44]. The contribution of adult neurogenesis to odor discrimination learning may then mostly result from the addition of new GCs that have enhanced plasticity while they are young, This includes synaptic-weight plasticity [45, 46] as well as structural spine plasticity, particularly during active learning [47], both of which guide the abGCs to connect to the appropriate MCs and top-down projections.

The MC-GC network is known to drive the *γ*-rhythms that are very prominent in the olfactory bulb [48]. They represent the collective dynamics of many neurons. It has been hypothesized that *γ*-rhythms may play an important role in the communication between brain areas [49, 50]. In the olfactory bulb, their power is enhanced with task demand after learning [51] and optogenetic excitation of GCs in the *γ*-range enhances discrimination learning [52], suggesting a role of *γ*-rhythms in odor processing. A natural outcome of our model are subnetworks comprised of interconnected MCs and GCs that respond preferentially to the same odors. When driven by the corresponding odors, each of these subnetworks is likely to generate its own *γ*-rhythm with its own frequency. The question then arises whether these rhythms will synchronize if the bulbar network is driven by the corresponding odor mixtures. Depending on the temporal window over which the bulbar output is read out in downstream brain areas, the perception of such mixtures may well vary with the degree of synchrony. Interestingly, the synchrony of such interacting rhythms can be enhanced by uncorrelated noise and by neuronal heterogeneity *via* a paradoxical phase response [53, 54].

A prominent feature of the olfactory bulb is its extensive centrifugal input *via* top-down projections that target the GCs particularly. Our model for the structural spine plasticity suggests that the GC develop receptive fields for specific odors of the training set and are preferentially connected with MCs with similar receptive fields. Due to the reciprocal nature of the MC-GC synapses, this implies that the activation of such a GC inhibits specifically MCs with the same response profile. If the top-down connectivity was able to target specific GCs, top-down inputs could control bulbar processing by inhibiting very specific MCs. Computational modeling [14] suggests that the extensive adult neurogenesis of GCs observed in the bulb naturally leads to a network structure that allows just that. That model predicts that non-olfactory contexts that have been associated with a familiar odor allow to suppress that odor, enhancing the detection and discrimination of novel odors. The spine plasticity investigated here leads to a network structure that is very similar to the bulbar component of the network obtained in the bulbar-cortical neurogenesis model. It is therefore expected to support the specific control of bulbar processing by higher brain areas.

## 4 Methods

### 4.1 Generation of simplified stimuli and naturalistic stimuli

The naturalistic stimuli are based on glomerular activation data [27], which contains 2-*d* imaging z-scores of the glomeruli responses for a variety of odorants. In our simulations, we used carvone, citronellol (pre-training), ethylbenzene, heptanal (training, pure, and mixture). Since in the original data, not all pixel values were available for all odorants, we kept only those that were common to all of them. We then downsampled the resulting 2074 data points to 240 sample points ***S***^(*orig*)^.

The z-score data ***S***^(*orig*)^ do not include the baseline activity of the glomeruli. Thus, a z-core of 0, which represents the mean activity across the whole population, does not imply that the activity was not affected by the odor presentation. To obtain a rough calibration of the mean activity, we used the observation that around 60% of the MCs are activated by any given strong stimulus [26]. In the absence of other information, we used this as a guide to re-calibrate the 40% percentile z-score *S*^(40%)^ as 0. Then, we normalized the data by their maximum, resulting in

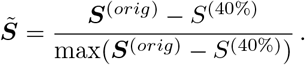

Further, MCs are activated by the airflow even without any odorants, and odor representations change mostly linearly with the concentration of odorants [55]. For any given mixture with concentration *p* of odor *A* and (1 − *p*) of odor *B* we therefore modeled the stimulus as

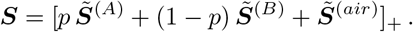

Here 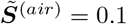 is the ‘air’ stimulus and [·]_+_ the rectifier, [*x*]_+_ = max(*x,* 0).

For the simplified model stimuli we used scaled Gaussian functions with different mean and standard deviation as 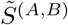 and set 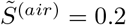.

### 4.2 Activation function

The neuronal network within the olfactory bulb is an excitatory-inhibitory network. To guarantee the firing rate is positive, the activation function *F*_*M,G*_ is required to be positive. Further, MCs within the olfactory bulb experience saturation behavior. In [55], the authors recorded responses of MCs when the concentration of odorant changed 10-fold. Among the recorded cells, 38% of MCs responded linearly to stimuli, and 29% of the MCs experienced saturation behavior. This saturation may be the result of saturation in the inputs to the MCs or in the MCs themselves. The activity of the sensory neurons saturates on a log scale when the odor concentration changes 1000-fold [56]. For the 10-fold concentration changes relevant in our studies, we therefore assumed that saturation occurred mostly in the MC rather than their sensory inputs and took the activation functions in (1,2) to be

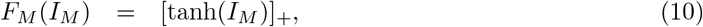

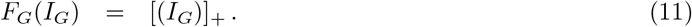

### 4.3 Stabilization Mechanism

For simplicity, we introduced top-*k* competition as a stabilization mechanism as follows. If a GC had more than *k* > *N*_*conn*_ synapses, only the top *k* synapses with largest *R*_*ij*_ survived to the next step. As a result, the number of synapses on each GC was bounded by *k*, while the initial number of synapses was *N*_*conn*_. A soft top-*k* competition can be realized by a resource-pool competition with additional assumptions, as discussed below. However, other mechanisms like scaling [57] or the ABS (Artola, Bröcher and Singer) rule [58] may alternatively restrict the total number of synapses. Since the stability mechanism was not our focus, we mostly used the (hard) top-*k* competition without directly specifying its biological cause.

### 4.4 Definition of responsive cells and divergent cells

We defined the response of an MC as the change in activity induced by an odor compared to the activity with air as stimulus. Specifically, if a MC had activity ***M*** ^(*A,B*)^ in response to stimuli ***S***^(*A,B*)^, and it had activity ***M*** ^(*air*)^ when presenting air as a stimulus ***S***^(*air*)^, we defined the response to odor *A, B* as 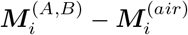.

The definitions of responsive and divergent cells were adapted from [16]. A cell was classified as responsive, if its response to one odor in an odor pair was above a given threshold *θ*, i.e. 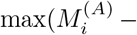 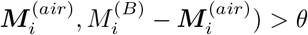. A cell was classified as divergent, if the difference in its response to the two odors was above the same threshold, 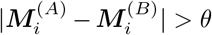. Our results were robust over a range of values of the threshold *θ* (Figure S12).

### 4.5 Characterizing Discriminability

We used d-prime and the Fisher discriminant to assess the discriminability of activity patterns. For each divergent neuron d-prime is given by the difference between the mean activities of the neuron for the two presented odors scaled by the combined variability of the activities. In our firing-rate model (1,2) the MC-activity exhibits fluctuations only on the time scale of network restructuring, which is much longer than the odor presentation. Assuming, as in [14], that the MC firing rates 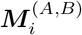 represent the mean values of independent Poisson spike trains, the variances **Σ**^(*A,B*)^ of those spike trains are then given by their respective means, 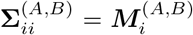 and 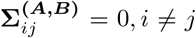. Thus, for each MC we have

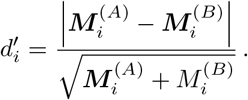

One way to characterize the overall discriminability of the activity patterns is then given by the average of the d-prime 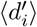 across all neurons, as was done in [16].

We also used the Fisher discriminant to measure the discriminability. In general, it is given by

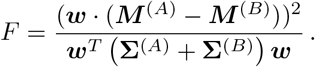

Here ***w*** is an arbitrary vector, which can be interpreted as the weights with which the MC activity is fed into a linear read-out neuron. The weights ***w*** that maximize *F* are given by 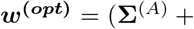 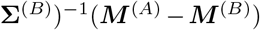. The optimal Fisher discriminant used to assess discriminability is then given by

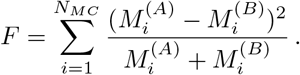

It is related to d-prime of the individual neurons *via*

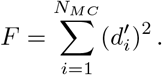

In contrast to 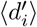, *F* increases with the number of MCs.

## 5 Acknowledgments

We thank S. Saha, K.A. Sailor, and P.-M. Lledo for discussions and for sharing their data with us prior to publication. We gratefully acknowledge discussions with T. Komiyama.

This work was supported by funding from the National Institutes of Health (https://www.nidcd.nih.gov/) through grant DC015137 to HR.

## 6 Supplementary Materials

### 6.1 Differential Inhibition and Disinhibition

**Figure S1:**
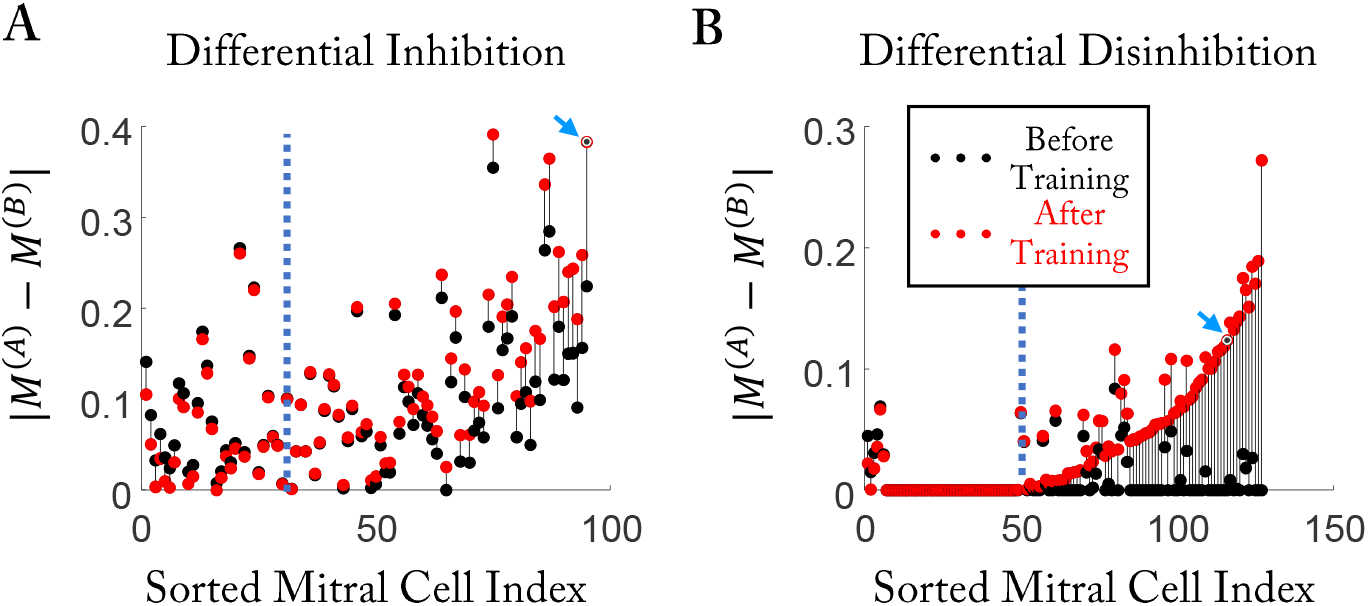
Differential behavior of MCs after training. Response differences 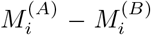 before (black) and after training (red). Cells are sorted by the change in the response difference from before to after the training. (A) MCs that are excited by both odors before training 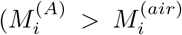, 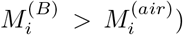. For the cells to the right of the dotted line differential inhibition enhanced the response difference. Blue arrow: cell shown in Figure 2J1. (B) MCs that are inhibited by both odors before training 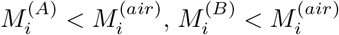. For the cells to the right of the dotted line differential disinhibition increases the response difference. Blue arrow: cell shown in Figure 2J2.

### 6.2 Robustness of the model

To demonstrate the robustness of the model and the influence of the various parameters on its performance, we assessed the impact of varying the major parameters of the model.

Changing the total number *N*_*GC*_ of GCs affected the performance very little, since the number of GCs that were strongly activated and connected to specific MCs did not change significantly, which influenced the effective connectivity *W* ^(*mm*)^ only mildly (Figure S2). Analogously, changes in the inhibitory strength *γ* were largely compensated by a change in the number of activated GCs (Figure S3). This feedback stabilizing the overall inhibition arises from the reciprocal character of the MC-GC connections: new connections increase the inhibition of the MCs, which decreases the excitation of the GC and in turn inhibits the formation of new connections.

What does control the overall inhibition, is the threshold *G*^(1)^ in the activation function *ϕ*(*G*) (Figure S4). Only GCs with activity larger than *G*^(1)^ form more synapses and generate stronger selective inhibition. Thus, for larger *G*^(1)^ fewer GCs had sufficiently large activity to sustain their synapses, which reduced the overall inhibition and increases the MC activity level. The change in overall inhibition could be associated with a change in the selectivity. To quantify this, we call a GC responsive to odor *A* (*B*), if after training its activity surpasses *G*^(1)^ for odor *A* (*B*). Despite the increase in the inhibition, the number of GCs that responded to both odors remained low when decreasing *G*^(1)^ (Figure S4H, red line), indicating that the selectivity depends very little on *G*^(1)^.

The other threshold in the activation function, *G*^(0)^, affects the removal but not the formation of synapses. For low values of *G*^(0)^ weakly activated GCs removed many of their synapses, particularly those connecting them to active MCs (Figure S5C, D). This lead to higher selectivity of the connectivity and with it to a weaker inhibition of the spontaneously active MCs that were not odor-driven (Figure S5B, E). It had, however, little impact on the maximal MC amplitudes (Figure S5H). The removal of synapses played a larger role in retaining and forgetting connections. To assess its impact, we trained the model in a second phase with a new pair of stimuli that overlapped with the stimuli learned in phase 1 (Figure S5I). For *G*^(0)^ = 0 any weak activity (0 < *G* < *G*^(1)^) triggered the removal of spines and lead in phase 2 to an almost complete forgetting of the connectivity learned in phase 1 (Figure S5D, K) and with it to poor discrimination of the previously learned odors (Figure S5N). However, for large *G*^(0)^ the previously learned connectivity was maintained (Figure S5G, M) and with it the discriminability of the previously learned stimuli. The discrimination of the newly learned stimulus pair 2 was, however, not significantly affected by *G*^(0)^ (Figure S5O).

**Figure S2:**
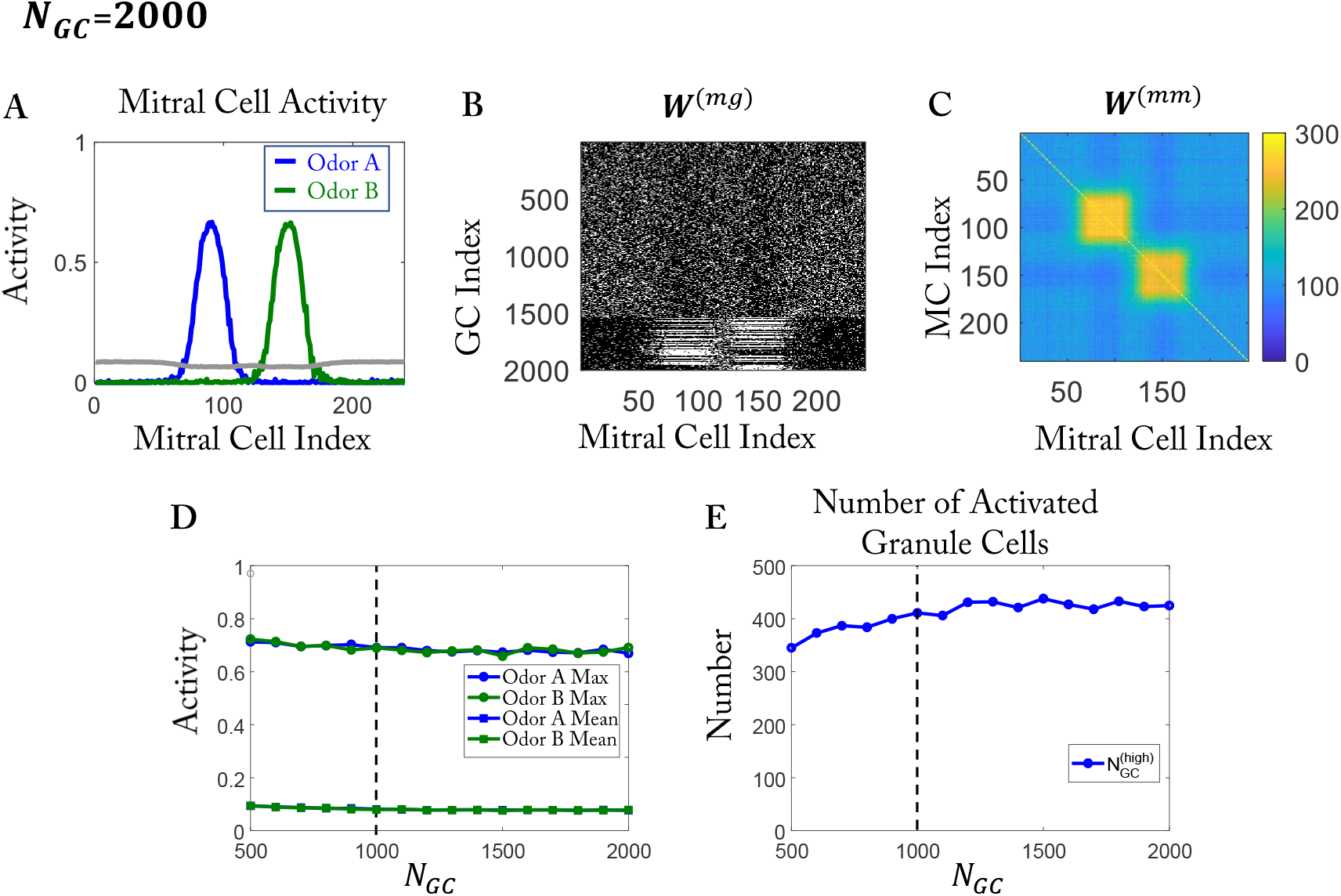
The overall inhibition of the model depends only weakly on the number *N*_*GC*_ of GCs. (A-C) Training with simplified easy stimuli as in Figure 1D and random initial network as in Figure 1F, but with twice as many GCs. (A) MC activity after training (cf. Figure 1H). (B) Connectivity *W* ^(*mg*)^ after training. (C) Effective connectivity *W* ^(*mm*)^. Compared to Figure 1J the connectivity is not quite as selective. (D) The maximal and mean MC activity after training as a function of the number of GCs. (E) The number of activated GCs (*G* > *G*^(1)^) depends only weakly on the total number of GCs. In (D, E) the vertical dashed line marks the value used in the rest of the paper.

The maximal number of connections *k* that each GC can make had a significant impact on MC amplitudes and on the selectivity of the connections. With increasing *k*, each GC integrated input from more MCs, and more easily surpassed the threshold *G*^(1)^. Each of these more activated GCs inhibited a larger number of MCs, increasing the overall inhibition (Figure S6C). When the maximal number of connections was larger than the number of MCs activated by one of the odors, not all connections of a GC could be made to MCs that were activated by odor *A*, say. The remaining connections could then be made to MCs corresponding to odor *B* without destabilizing the other synapses. As a result, GCs that responded to both odors emerged (Figure S6D).

**Figure S3:**
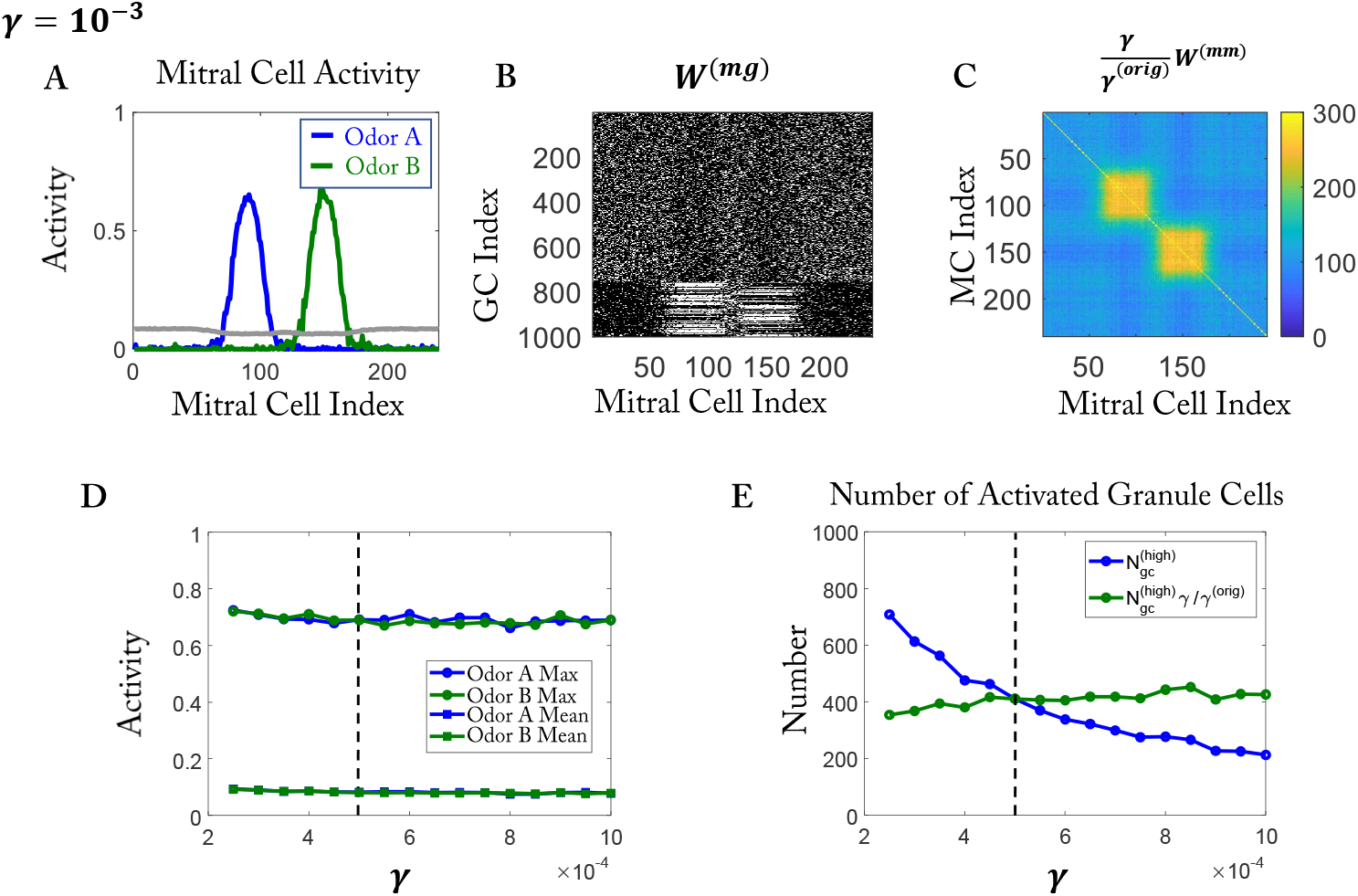
The overall inhibition of the model depends only weakly on the inhibitory strength *γ*. The results are organized as in Figure S2. (C) The connectivity is slightly less selective than in Figure 1J. (D,E) The vertical dashed line indicates the value used in the rest of the paper. (D, E) To allow a direct comparison, *W* ^(*mm*)^ and 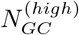 have been re-scaled by 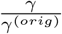 with *γ*^(*orig*)^ = 5 × 10^−4^.

To further illustrate that the network learned not by simply changing the number of synapses, but by developing a specific, stimulus-dependent connectivity, we assessed the performance of the network when the connections of each GC were rewired to random MCs with probability *p*_*rewire*_ while keeping the number of synapses on each GC the same. Fr *p*_*rewire*_ = 0 (Figure S7A), the connection was the originally learned network; for *p*_*rewire*_ = 1, the connection for each GC was totally random (Figure S7C). Indeed, for the highly similar training odors, the discriminability predominantly decreased as the randomness is increased (Figure S7D). For the network trained on the dissimilar odors (easy task), the discriminability increased, as expected, when the randomness was increased (Figure S7F). Interestingly, when *P*_*rewire*_ was increased beyond 0.7 these trends reversed (Figure S7D, F) somewhat.

The task of distinguishing odors can be hard in different ways. Motivated by the experimental results in [16], we considered so far only similar mixtures comprised of dissimilar odors. These stimuli differed most in the most activated MCs. However, a task can be hard because the stimuli differ only in the weakly activated MCs and are identical in the strongly activated ones. This would arise, for instance, if two odors need to be distinguished in the presence of a common, strong background odor. We therefore trained and tested the model with stimuli in the form of skewed Gaussians, which differed predominantly in their weak components (Figure S8A). Again, the training enhanced the response differences for many MCs (Figure S8E) and increased the Fisher discriminant (Figure S8F).

**Figure S4:**
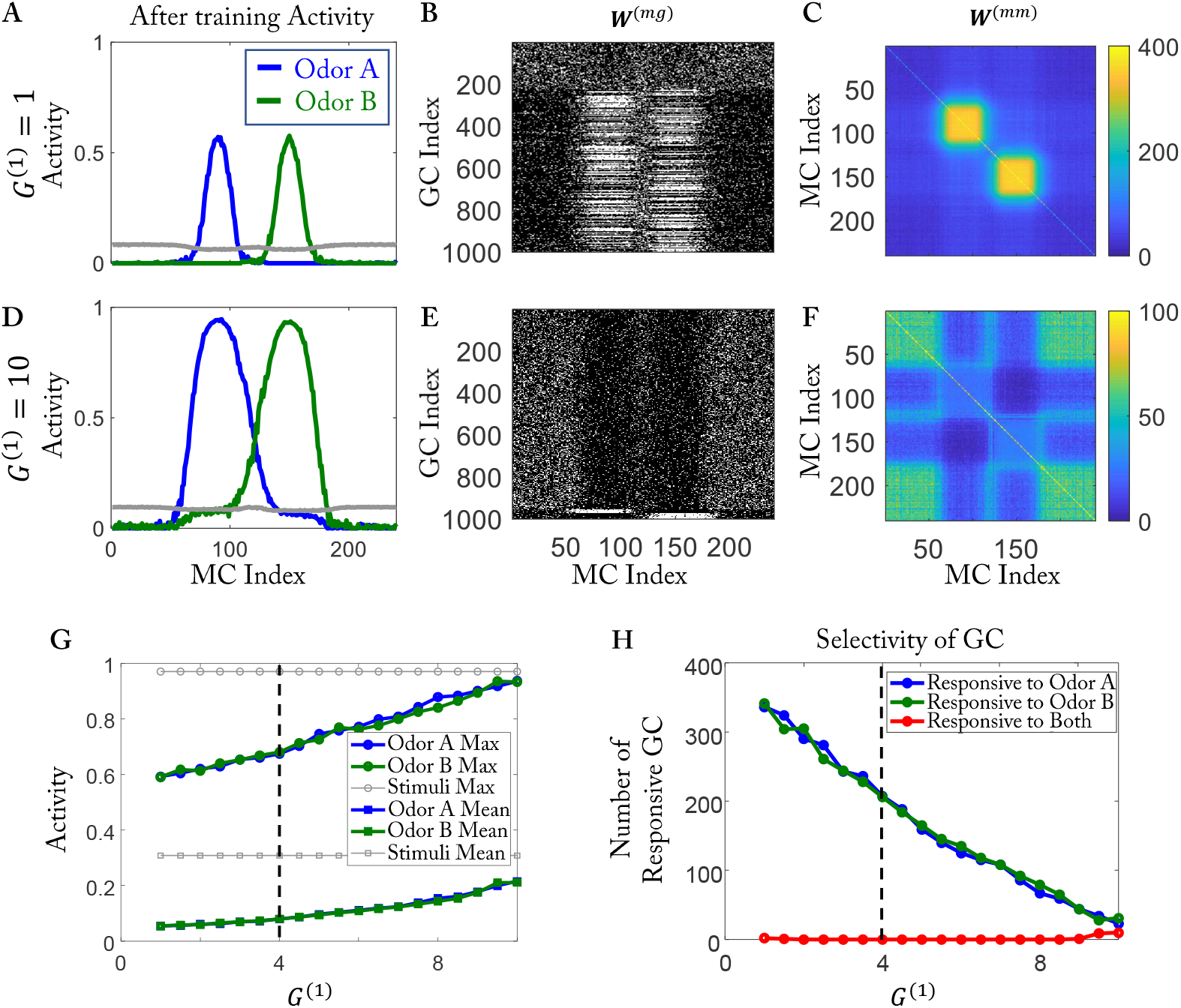
Increasing *G*^(1)^ impairs the ability to form connections between activated MCs and GCs. (A,B,C) Results for *G*^(1)^ = 1. (A) MC activity after training. (B) Connectivity *W* ^(*mg*)^ between MCs and GCs. (C) Effective connectivity *W* ^(*mm*)^. (D,E,G) as (A,B,C) except for *G*^(1)^ = 10. Activated MCs are connected with fewer GCs (compare E with B), resulting in weaker disynaptic inhibition (compare F with C) and higher MC activity (compare D to A). In (F) most of the GCs cannot reach the high threshold. As a result, the synapses that connect to strongly activated MCs are removed faster than those connecting to weakly activated MCs. Thus, the effective connectivity among the activated MCs is lower than the background. (G) Maximal and mean MC activity increase with increasing *G*^(1)^. The gray lines indicate the corresponding values without inhibition from GCs (by setting *γ* = 0). (H) The number of responsive GC decreases with increasing *G*^(1)^. (G,H) The vertical dashed line indicates the value used in the rest of the paper.

**Figure S5:**
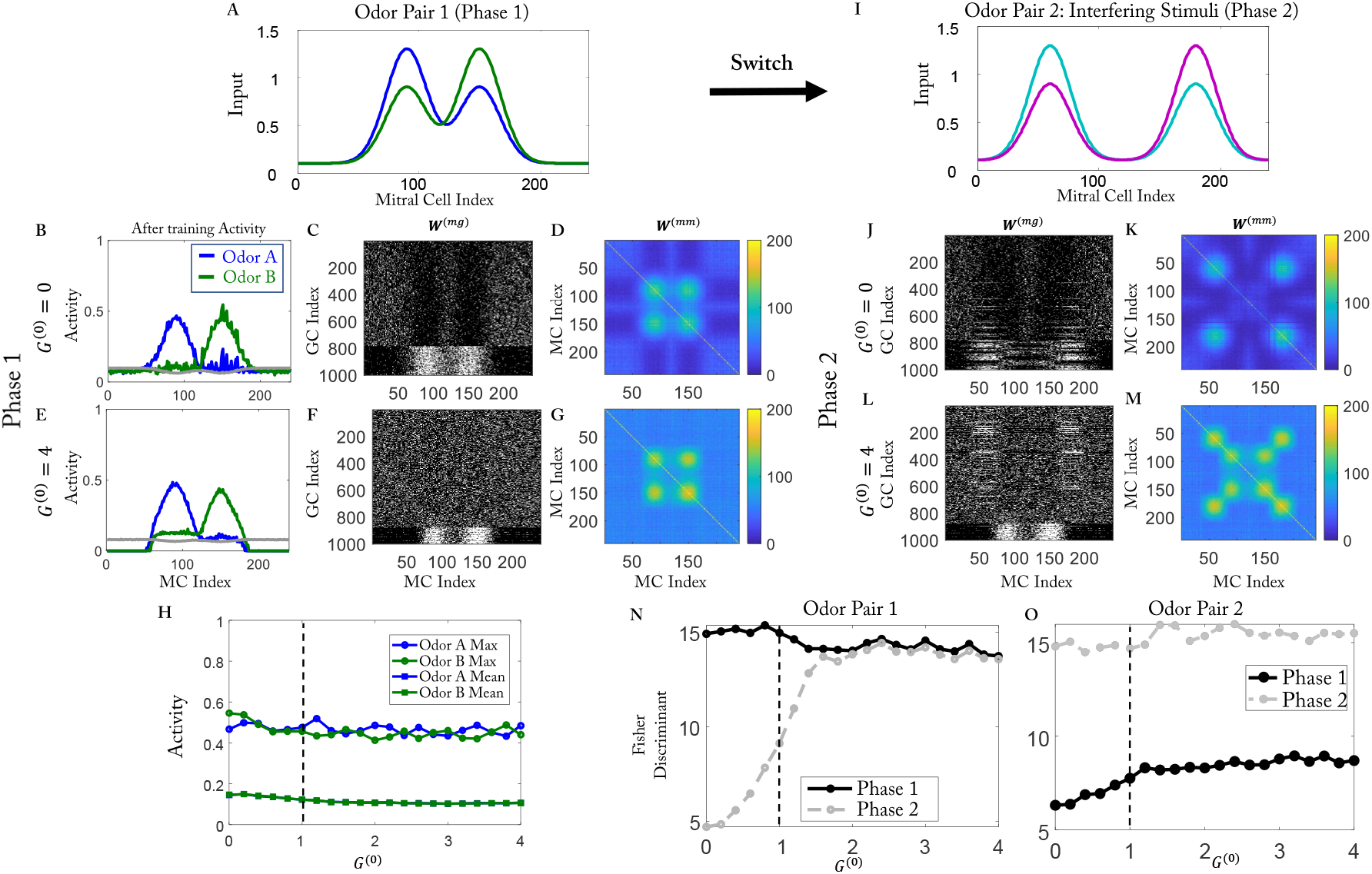
Learning ability is not affected by *G*^(0)^, but the memory is. (A-G) Phase 1. (A) Training stimuli in phase 1. (B) MC activity after training. (C) Connectivity *W* ^(*mg*)^. (D) Effective connectivity *W* ^(*mm*)^. (E to G) as (B to D) except for *G*^(0)^ = 4. (H) Maximal and mean MC activity after training in phase 1 as a function of *G*^(1)^. (I to M) as (A to G) but after Phase 2. (I) Interfering training stimuli. (J-M) For *G*^(0)^ = 0 the network forgets the previously learned connectivity, but not for *G*^(0)^ = 4. (N) The Fisher discriminant for odor pair 1 is unaffected by *G*^(1)^ in phase 1, but substantially reduced in phase 2 for small *G*^(1)^. (O) For odor pair 2 the Fisher discriminant depends only little on *G*^(1)^. The vertical dashed line in (H,N,O) indicates the value used in the rest of the paper.

**Figure S6:**
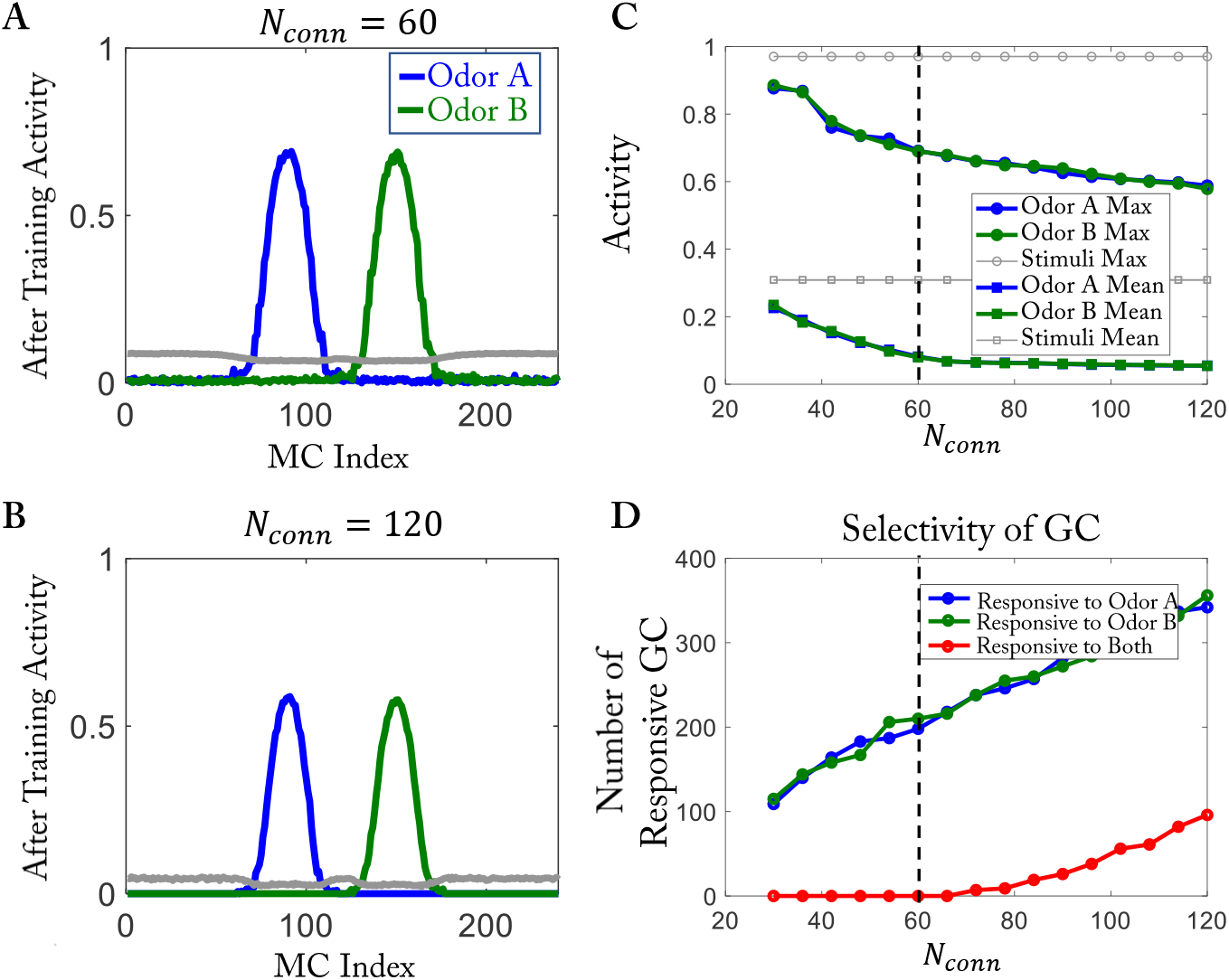
The selectivity of GCs is influenced by the number of connections each GC makes. Easy stimuli as in Figure 1D with random initial network as in Figure 1F. The ratio of maximal to initial number of connections is kept fixed at *k/N*_*conn*_ ≡ 1.1. (A, B) The MC activity is very similar for *N*_*conn*_ = 60 and *N*_*conn*_ = 120. (C) Maximal and mean MC activity decrease with increasing *N*_*conn*_. The gray lines indicate the corresponding values without inhibition from GCs (by setting *γ* = 0). (D) The selectivity of the GCs is impaired for *N*_conn_ > 60: a population of cells emerges that respond to both odors (red line).

**Figure S7:**
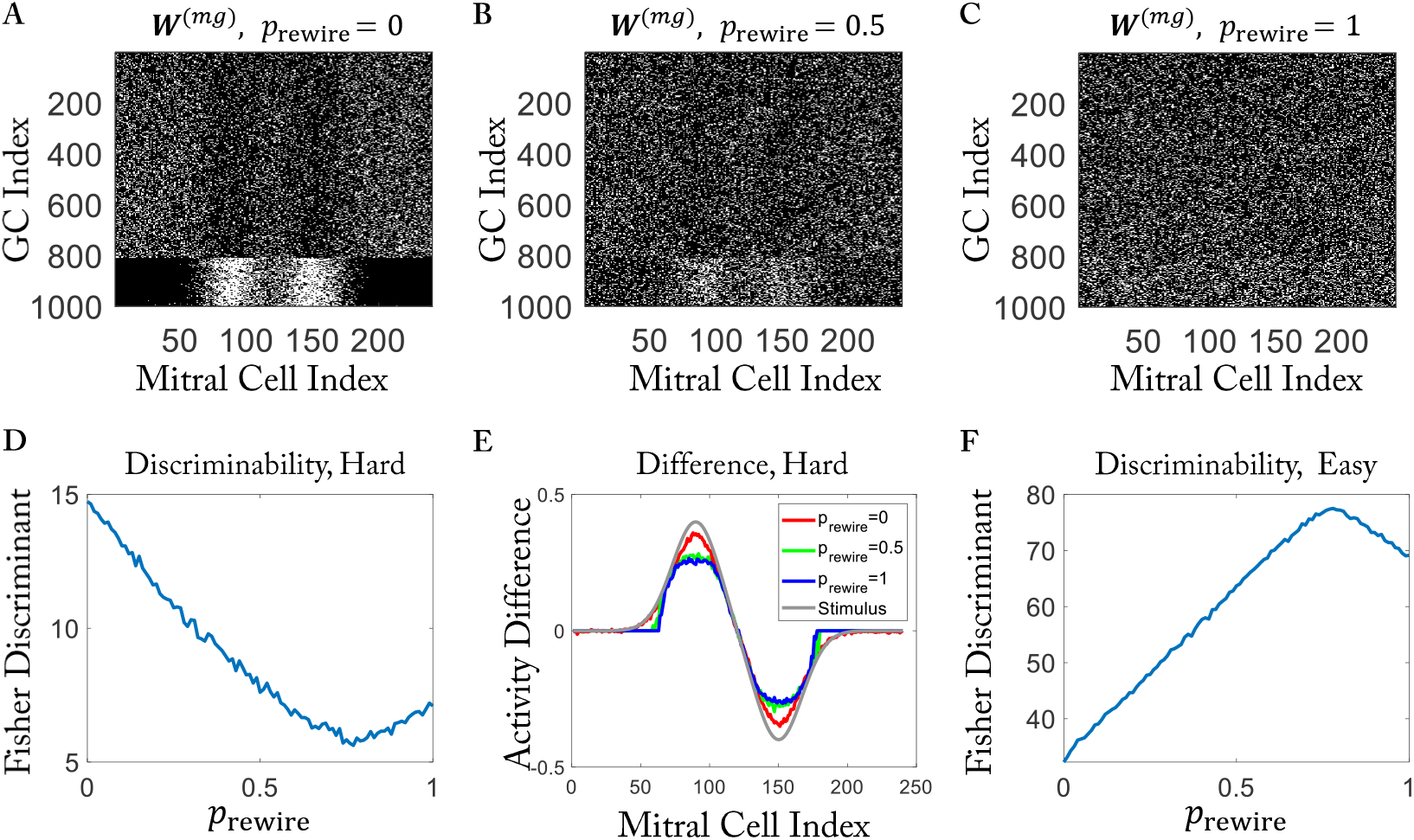
Rewiring trained networks. (A to C) Connectivity matrices after rewiring a network that was trained with the hard task (Figure 3C, bottom). Rewiring probability *p*_*rewire*_ = 0, 0.5, 1, respectively. (D) For the hard task discriminability predominantly decreases with increasing rewiring probability *p*_*rewire*_. (E) Activity differences are reduced after rewiring. (F) For the easy task discriminability predominantly increases with increasing *p*_*rewire*_ (cf. Figure 3C, top).

**Figure S8:**
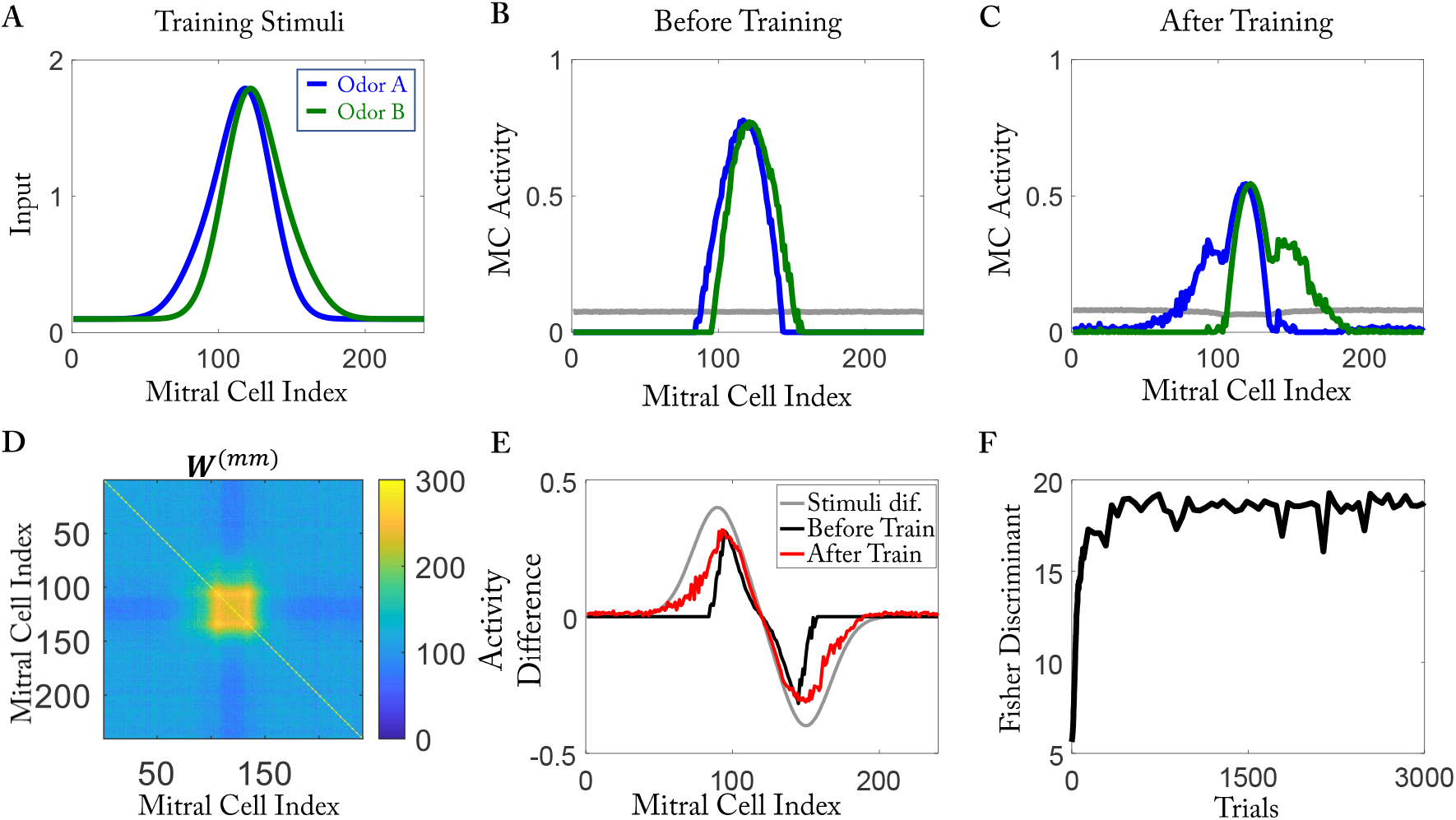
Discrimination of stimuli differing mostly in weakly activated MCs. (A) Training stimuli (cf. Fig.3. (B, C) MC activity before and after training. (D) Effective connectivity matrix *W* ^(*mm*)^ after training. (E) Activity differences are increased by training. (F) The Fisher discriminant increases with training. In order to suppress noise, the number of GC and the simulation were doubled (*N*_*GC*_ = 2000).

**Figure S9:**
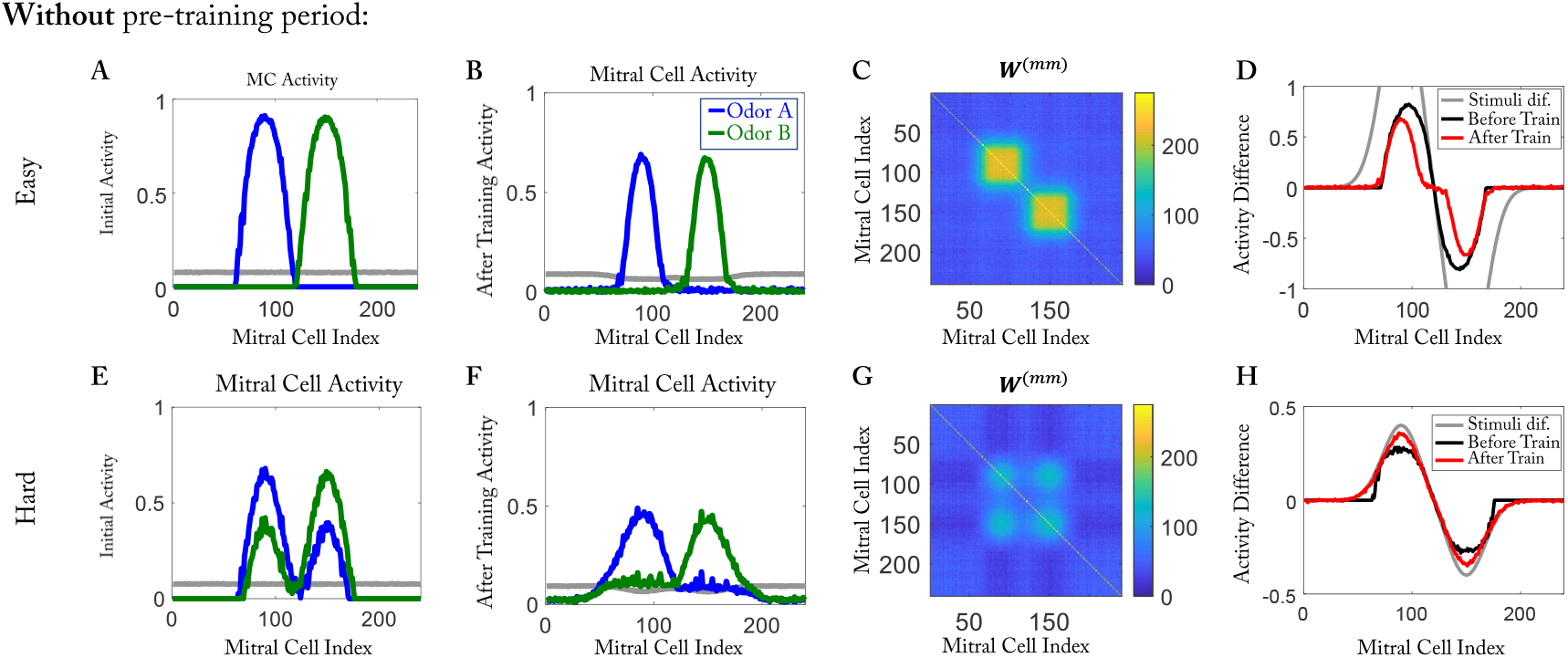
Training results without pre-training phase. The training starts with a random homogeneous network. Results are organized as in Figure 3E to L.

**Figure S10:**
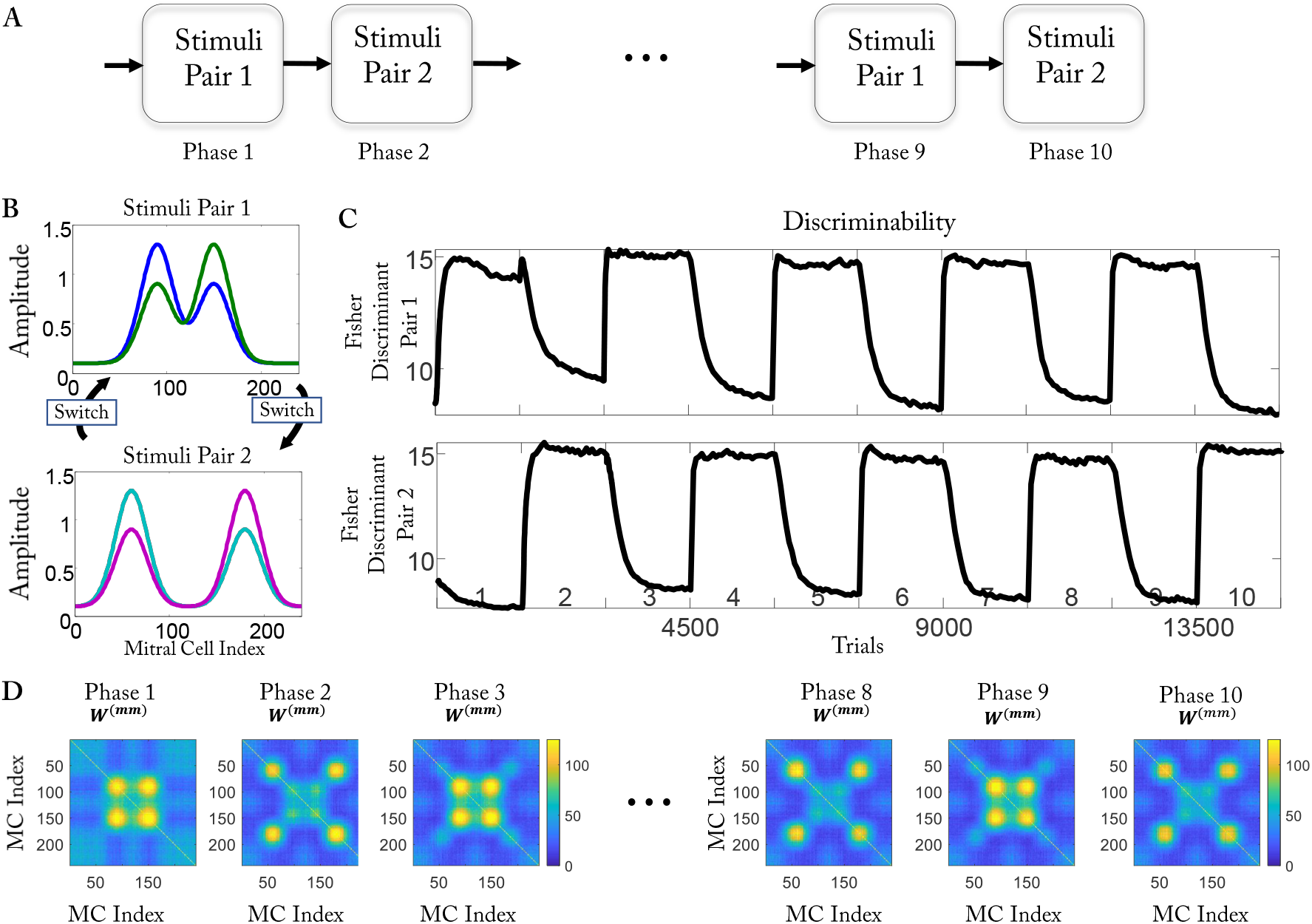
Alternating training does not impair learning ability. (A) Expanding the training protocol of 4A to 10 phases. (B) (Top) Stimuli of odor pair 1. (Bottom) Interfering stimuli of odor pair 2. (C) The Fisher discriminant of odor pair 1 (top) and pair 2 (bottom). Learning (increasing Fisher discriminant) proceeds faster than forgetting (decreasing Fisher discriminant). The numbers above the x-axis indicate the phase. (D) Effective connectivity *W* ^(*mm*)^ alternates.

**Figure S11:**
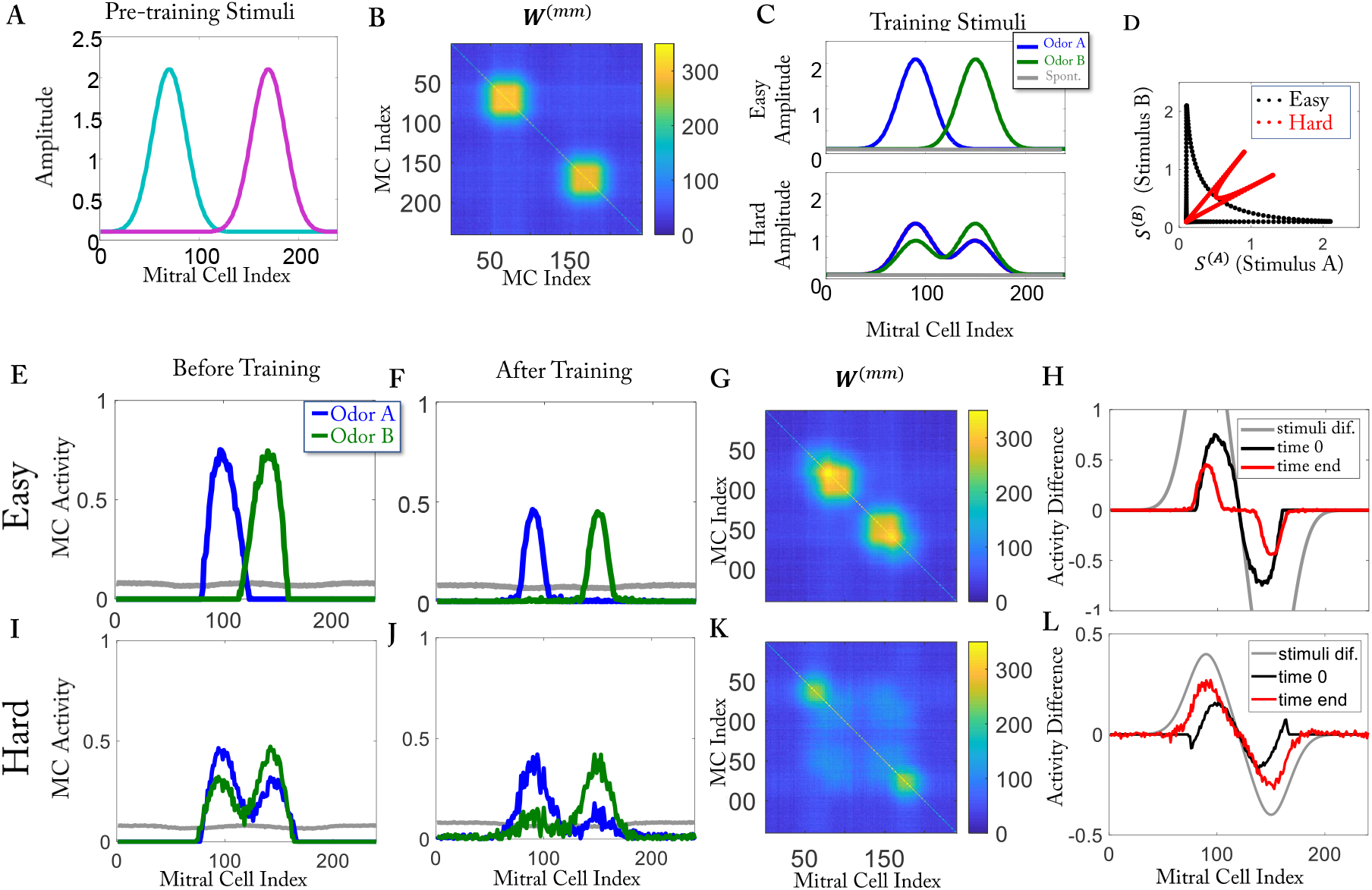
Results of the resource-pool model when trained with simplified stimuli. Here *γ* = 1.7e 3. Results are organized as in Figure 3.

**Figure S12:**
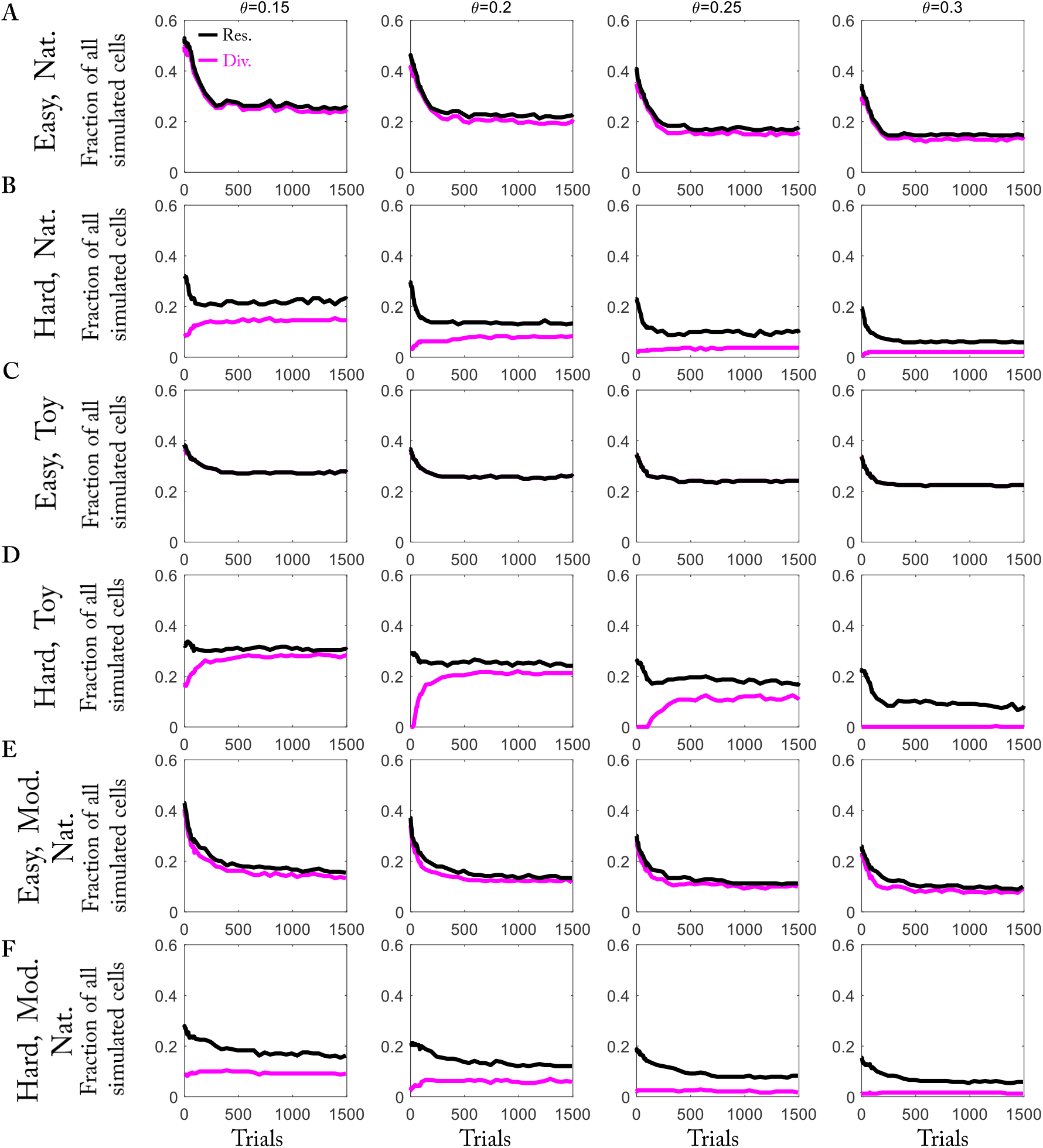
Qualitatively, the results for the fraction of responsive and divergent cells do not depend on the threshold *θ* for their classification. (A, B) Training with naturalistic stimuli for different thresholds *θ*. The results in Figure 2 and Figure 5 are based on *θ* = 0.2. (C, D) as (A, B) but for training with simplified stimuli. (E, F) as (A, B) but in the resource-pool model.

## Notes

### Competing Interest Statement

The authors have declared no competing interest.

